# Theta oscillations and minor hallucinations in Parkinson’s disease reveal decrease in frontal lobe functions and later cognitive decline

**DOI:** 10.1101/2022.11.24.517668

**Authors:** Fosco Bernasconi, Javier Pagonabarraga, Helena Bejr-Kasem, Saul Martinez-Horta, Jaime Kulisevsky, Olaf Blanke

## Abstract

Cognitive decline and hallucinations are common and debilitating non-motor symptoms, occurring during later phases of Parkinson’s disease (PD). Minor hallucinations (MH), appear at early phases and have been suggested to predict cognitive impairment in PD, however, this has not been well-established by clinical research. Here, we investigated whether non-demented PD patients with MH show altered brain oscillations and whether such MH-related electrophysiological changes are associated with cognitive impairments that increase over time. Combining model-driven EEG analysis with neuropsychiatric and neuropsychological examinations in 75 PD patients, we reveal enhanced frontal theta oscillations in PD patients suffering from MH and link these oscillatory changes with lower cognitive frontal-subcortical functions. Neuropsychological follow-up examinations five years later confirmed MH-specific theta oscillations and revealed a stronger decline in frontal-subcortical functions in MH-patients with stronger frontal theta alterations, defining an MH and theta oscillation-based early marker of a cognitive decline in PD.

## Introduction

PD affects approximately 3% of the population over 65 years (Kalia & Lang, 2015) and the number of patients is expected to double by 2040, reaching an estimated total of 15-18 million people worldwide and rising faster than any other neurological disorder (Dorsey et al., 2018). Although PD is traditionally defined as a movement disorder with the typical motor symptoms of resting tremor, rigidity, and bradykinesia, PD pathology also affects several non-motor circuits leading to a wide variety of non-motor symptoms that appear early in the disease course (Postuma & Berg, 2016). Among the latter symptoms, hallucinations are highly prevalent (Fénelon et al., 2010; Lenka et al., 2019), with one individual out of two experiencing hallucinations regularly (Ffytche et al., 2017; Lenka et al., 2019), increasing up to 70% at advanced stages of the disease (Aarsland et al., 2021; Levin et al., 2016), and often becoming a dominant non-motor symptom, together with dementia (Ffytche et al., 2017). Hallucinations in PD have major negative impact on patients, families, and society, and have been associated with a more severe form of the disease with poor cognitive outcome and dementia (Aarsland et al., 2003; Anang et al., 2014; Diederich et al., 2009; Fénelon et al., 2010; Forsaa et al., 2010; Galvin et al., 2006; Lenka et al., 2019; Marinus et al., 2018; Uc et al., 2009), as well as earlier home placement (Aarsland et al., 2000; Diederich et al., 2009; Goetz et al., 2008; Gonzalez et al., 2022; Marinus et al., 2018), and higher risk of mortality (Aarsland et al., 2000; Gonzalez et al., 2022).

Clinical and neuroimaging studies, investigating the pathophysiology of hallucinations in PD (Collerton et al., 2005; Goldman et al., 2014; Hobson et al., 2000; Ibarretxe-Bilbao et al., 2010; Shine et al., 2011; Watanabe et al., 2013), consistently highlighted impairments in posterior visual-perceptual and frontal executive-attentional functions and related brain networks. Impairments in visual-perceptual functions are consistent with the frequent visual nature of hallucinations in PD (formed visual hallucinations, VH) (Goldman et al., 2014; Ramirez-Ruiz et al., 2007; Watanabe et al., 2013) and alterations in fronto-striatal networks have been proposed to account for executive frontal-subcortical deficits (Obeso et al., 2014; Shine et al., 2014). Both findings link the occurrence of hallucinations with cognitive impairments and related brain networks (Aarsland et al., 2003; Anang et al., 2014; Galvin et al., 2006; Uc et al., 2009). This past work on hallucinations has predominantly focused on VH, that is on structured complex VH of people or animals. However, because VH occur most often at later stages of the disease, with cognitive decline already present, they are not suitable as an early marker of cognitive decline in PD, as has been proposed for minor hallucinations (MH).

MH consist of presence hallucinations, passage hallucinations, and pareidolias (Fénelon et al., 2000; Ravina et al., 2007). MH are usually experienced at earlier stages of the disease (Ffytche et al., 2017; Lenka et al., 2019) and can even precede parkinsonian motor symptoms (Pagonabarraga et al., 2014). However, compared to VH, much less is known about the brain mechanisms of MH, although they have recently been shown to involve fronto-temporal brain regions related to executive sensorimotor function (Bernasconi et al., 2021). Brain alterations in patients with MH have also been reported to partially overlap with brain regions identified for VH (Bejr-Kasem et al., 2019; Pagonabarraga et al., 2014) and for cognitive deficits in PD (Bejr-Kasem et al., 2021; Bernasconi et al., 2021). Despite these promising findings, previous clinical work did not reveal cognitive impairments in PD patients with MH (versus those without MH), in either visual-perceptual and/or executive frontal-subcortical functions (Bernasconi et al., 2021; Bejr-Kasem et al., 2021; Llebaria et al., 2010; Pagonabarraga et al., 2014), suggesting that the neuropsychological assessment instruments alone are not sensitive enough to detect early alterations associated with MH and that more sensitive measures are needed. Furthermore, it is currently not known whether MH could contribute to the early detection of PD patients at risk of developing cognitive impairment.

Beside neuropsychological and psychiatric investigations, electroencephalography (EEG) has also been explored in cognitive decline assessment in many neurological disorders, including PD (e.g. Bosboom et al., 2006; Geraedts et al., 2018; Hassan et al., 2017; Olde Dubbelink et al., 2013). EEG recordings have the advantage to be widely available (in most hospitals, clinics, and even at patient homes; Hinrichs et al., 2020), and allow for non-invasive measurements of brain activity (e.g. oscillations). In PD, analysis of neural oscillations has revealed motor-related changes, such as enhanced beta oscillations in subcortical regions (e.g. subthalamic nucleus) and between subcortical structures and the motor cortex (e.g. Hirschmann et al., 2011, 2013; Kühn et al., 2009; Oswal et al., 2016; Tinkhauser et al., 2017). Concerning cognitive decline in PD, cross-sectional studies comparing patients at different stages of the disease (e.g. PD patients with normal cognition vs. PD with cognitive impairments, independent from hallucinations) revealed enhanced power in lower frequency bands, as well as a reduction in higher frequency bands (e.g. Babiloni et al., 2020; Bonanni et al., 2008; Bosboom et al., 2006; Geraedts et al., 2018; Hassan et al., 2017; Olde Dubbelink et al., 2013). Despite the putative role of such latter oscillatory changes in cognitive decline, it is not known whether these changes can already be detected in patients with early PD (with no or mild cognitive impairments) or whether they are only manifest at later stages of the disease, associated with prominent cognitive impairments. Critically for the present study, it is currently unknown how the presence of MH is related to brain oscillations and whether the blending of neuropsychological, neuropsychiatric, and EEG data might help in detecting subtle cognitive changes in PD patients with MH, at an early stage of the disease.

Here, we investigated whether PD patients with MH show altered brain oscillations and whether such MH-related electrophysiological changes are associated with an early neuropsychological impairment that increases over time. In 75 PD patients, we applied a model-driven EEG approach (Donoghue et al., 2020) to measure periodic and aperiodic properties of the resting-state EEG data and combined it with comprehensive neuropsychiatric interviews (to determine MH and VH) and with neuropsychological examinations (to measure cognitive function). We show that the neuropsychological performance of patients with MH is comparable to those without MH, confirming earlier data that neuropsychology alone is not sensitive enough to detect early alterations associated with MH. However, PD patients with MH showed oscillatory alterations in the theta band over frontal regions, which were MH-specific. Critically, these MH-specific changes in theta oscillations were associated with lower cognitive frontal-subcortical functions, allowing us to define an MH and theta oscillation-based marker of a subclinical cognitive impairment. Finally, to assess cognitive functions over time and their relation to the present MH-specific theta marker we conducted neuropsychological follow-up examinations at two years (available in 68 patients) and at 5 years (available in 54 patients) after the first assessment. These follow-up data confirmed MH-specific theta oscillations and, critically, revealed that the decline in frontal-subcortical functions is more rapid and more severe in MH-patients with stronger frontal theta alterations, as measured five years earlier.

## Results

### Minor hallucinations (first assessment, semi-structured interview)

Based on the neuropsychiatric semi-structured interview, 75 patients with PD were grouped into those who reported MH (PD-MH; n = 31) and those without MH (PD-nMH; n = 44). In the PD-MH subgroup, in addition to MH, two patients also reported VH, and one patient reported auditory hallucinations. Although the exact prevalence of MH in PD is still debated, evidence from previous studies suggests that approximately 50% of patients have MH, with presence and passage hallucinations as most the frequent MH (Fénelon et al., 2000, 2011; Pagonabarraga et al., 2016; Wood et al., 2015). Our results corroborate the prevalence of MH in PD, with 41.37% (31/75) of the patients experiencing MH. Patients can experience one MH only (e.g. presence hallucination) or multiple MH (e.g. presence and passage hallucination, not necessarily concomitant) (Figure 1).

**Figure 1.**
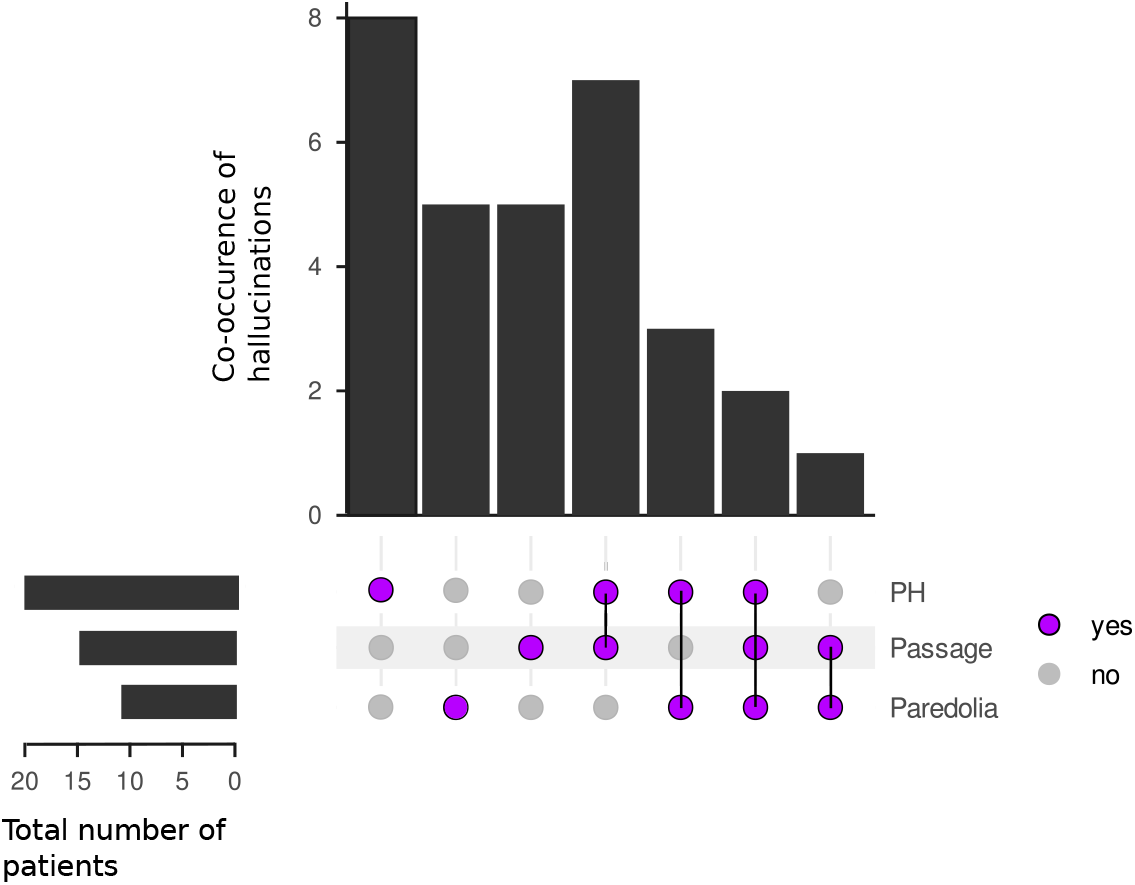
Prevalence of MH in PD. Prevalence of MH in PD. The figure illustrates the sum of patients with a specific MH (left bar plot) among the PD-MH. The figure also illustrates the sum of patients experiencing only one MH (violet dots indicate which MH, top bar plot indicates the sum of patients experiencing the MH) as well as the co-occurrence of two MH (two or more violet dots linked by a black line indicate which MH are co-experienced, the top bar plot indicates the sum of patients experiencing those MH). The figure was generated with UpSetR (Conway et al., n.d.) an R package.

### Demographical, clinical, and neuropsychological data (first assessment)

Our demographic data did not show any significant (all p-values > 0.05) differences in age, disease duration, or gender between the two patient subgroups (see Table 1 for more details). MH have been associated with the dosage of dopaminergic medications (e.g. Kataoka & Ueno, 2015). Yet, they have also been observed before the onset of motor symptoms and in the absence of any intake of dopaminergic medications (Pagonabarraga et al., 2014), leaving the role of those treatments in MH unresolved. Our results show that the dosage of the dopaminergic treatments were not significantly (p-value = 0.3 and p-value = 0.9; for levodopa equivalent dose and dopamine agonists equivalent dose, respectively) different between PD-MH and PD-nMH, supporting the hypothesis that MH are linked to PD rather than medication-related side effect (Ffytche et al., 2017, 2017; Pagonabarraga et al., 2016).

**Table 1.** Clinical and demographic variables for PD-MH and PD-nMH. Appendix *a* indicates Welch test, *b* Chi-squared, and *c* indicates U Mann Whitney test.

|  | PD-MH (N = 31) | PD-nMH (N = 44) | p-values |
| --- | --- | --- | --- |
| Age (years) | 67.9 ± 7.31 | 66.1 ± 8.63 | 0.34<sup>a</sup> |
| Gender (M/F) | 23/8 | 26/18 | 0.27<sup>b</sup> |
| Disease duration (years) | 5.74 ± 2.22 | 4.77 ± 2.11 | 0.06<sup>a</sup> |
| Education (years) | 12.7 ± 5.04 | 12.2 ± 4.68 | 0.7<sup>a</sup> |
| Eq. dopamine agonists (mg) | 162 ± 137 | 153 ± 99.4 | 0.74<sup>a</sup> |
| LEDD (mg) | 534 ± 250 | 464 ± 210 | 0.25<sup>a</sup> |
| UPDRS-III ON | 26.5 ± 8.75 | 25.2 ± 7.30 | 0.49<sup>a</sup> |
| PD-CRS Frontal-subcortical | 60.2 ± 15.4 | 63.8 ± 15.1 | 0.32<sup>a</sup> |
| PD-CRS Posterior | 28.5 ± 1.95 | 28.2 ± 1.94 | 0.42<sup>a</sup> |
| Hoehn & Yahr stage | 2.33 ± 0.38 | 2.18 ± 0.3 | 0.08<sup>c</sup> |

The results of the neuropsychological examination did not reveal any significant differences in cognitive functions between the PD-MH and PD-nMH patients. This was neither found for the frontal-subcortical cognitive functions (p-value = 0.3) nor for the posterior functions (p-value = 0.2). The two patient subgroups also did not differ (X^2^ (1,73) =1.33; p-value = 0.25) in the number of patients with mild cognitive impairment (MCI; defined as a PD-CRS total score ≤ 81 (Fernández de Bobadilla et al., 2013)), PD-MH (yes/no MCI): 10/21; PD-nMH (yes/no MCI): 9/35. These results are in line with previous literature showing that, at the group level, PD patients with MH do not differ from those without MH (Pagonabarraga et al., 2016) in cognitive functions. We also analyzed whether the number of MH (sum of different hallucinations) experienced by a patient was associated with the neuropsychological scores. Again, neither the frontal-subcortical cognitive score (rho = -0.14, p-value = 0.46) nor the posterior cognitive score (rho = 0.04, p-value = 0.84) was associated with the number of MH. These data show that PD-MH and PD-nMH do not differ in the demographic, clinical, and neuropsychological variables. The only difference between the two groups is the occurrence of MH. In addition, the amount the MH is not directly associated with cognitive impairment.

### Periodic and aperiodic EEG signals

The EEG signal is characterized by the background activity (1/f aperiodic signal; characterized by offset and slope; Supplementary Figure S1) and the genuine oscillatory signals (periodic signals; characterized by the center, power, and bandwidth of the peak; Supplementary Figure S1). Here we separated the aperiodic from periodic signals to better understand and interpret EEG results (Donoghue et al., 2020; see methods), focusing on the power and center frequency of the dominant oscillation; our analysis of the aperiodic signal included both the slope and offset. Based on the goodness of fit metrics, data were well fitted and no differences in fitting were observed between the two sub-groups of patients (see Figure S2 and Supplementary results).

### Frontal theta power (first assessment) is associated with lower frontal-subcortical cognitive functions in PD-MH patients, but not in PD-nMH patients

We analyzed whether the oscillatory power is modulated by MH and whether the oscillatory power is associated with lower cognitive performance (frontal-subcortical and posterior PD-CRS). Our results show that the frontal oscillatory power within the theta frequency band (4-8Hz; Figure 2A) was significantly (p-values <0.05; FDR corrected; Supplementary Table S1) modulated by the frontal-subcortical cognitive functions and by MH (yes/no) (i.e. interaction between the two terms) (Figure 2B). Post-hoc analysis revealed that the association between theta power and frontal-subcortical cognitive functions was significant for PD-MH (p-value < 0.001; Figure 2C), but not for PD-nMH patients (p-value = 0.28; Figure 2D). Moreover, our results indicate that for PD-MH patients the theta oscillatory power was negatively associated with the frontal-subcortical score, with higher theta power associated with lower cognitive scores (Figure 2C).

**Figure 2.**
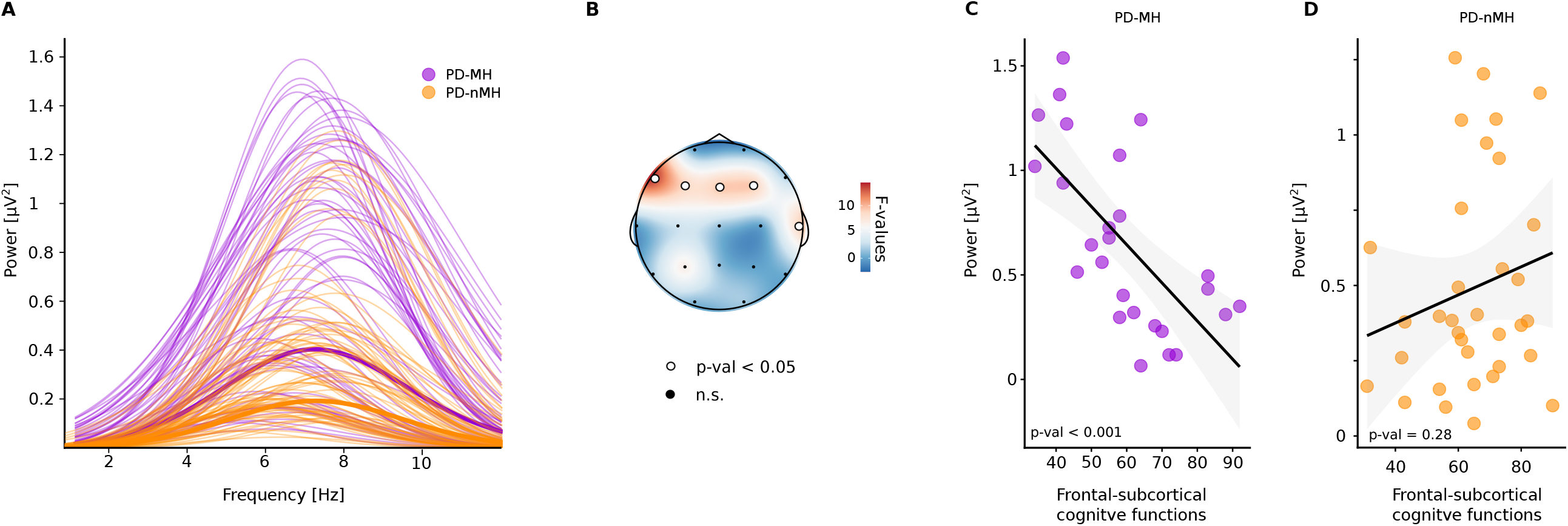
Association between frontal-subcortical cognitive functions and frontal theta power in PD-MH. **A**. Reconstructed aperiodic-adjusted theta peaks, for PD-MH (violet) and PD-nMH (orange), thicker lines indicate the mean of each group. Thinner lines indicate the single patient data. **B**. Topography of F-values indicating the interaction between patients’ group and frontal-subcortical cognitive functions. White highlighted dots indicate electrodes showing a significant (p-values < 0.05; FDR-corrected) interaction between the theta frequency band and frontal-subcortical cognitive functions. Note that 4 of 5 electrodes were over frontal scalp regions, bilaterally. **C**. Frontal theta oscillatory power is associated with frontal-subcortical cognitive functions in PD-MH. Higher power is associated with lower cognitive functions. Single dots represent the value for each patient (average of the electrodes showing a significant interaction between patients’ group and frontal-subcortical cognitive functions). **D**. Frontal theta oscillatory power is not associated with frontal-subcortical cognitive functions in PD-nMH. Single dots represent the value for each patient. Significance was obtained with permutation tests and multiple comparisons were corrected with FDR.

This association between oscillatory activity and frontal-subcortical functions (as a function of MH; interaction MH and frontal-subcortical PD-CRS) was specific for the theta frequency band, as no other frequency band (alpha: 8-13Hz, beta: 13-30Hz, and gamma: 31-45Hz) showed an association with frontal-subcortical cognitive functions (all interactions p-values > 0.05; FDR corrected; Supplementary Tables S1). In addition, this association was specific for electrodes over the frontal region, as no other electrode showed a significant (p-values > 0.05; FDR corrected; Supplementary Tables S1) for this interaction. Finally, the frontal theta pattern in PD-MH patients was only found for frontal-subcortical functions, as there was no association between MH and the posterior cognitive functions in any of the tested frequencies and electrodes (all interactions p-values > 0.05; FDR corrected; Supplementary Table S2). These data show that by merging neuropsychological, neuropsychiatric and EEG data, we are able to identify that selective signs of decreases in cognitive frontal-subcortical functions in PD are associated with MH and enhanced localized (frontal) theta power (see supplementary results for power changes associated with MH and Table S4-S5).

Because enhanced theta (and beta) power has been associated with motor impairment and its severity in PD patients (Asch et al., 2020; Beudel et al., 2017) and because the degree of motor impairment has been associated with cognitive impairments (e.g. Burn et al., 2006), it could be argued that the present findings (i.e. enhanced theta oscillations associated with MH and frontal-subcortical cognitive impairments) may reflect differences in motor symptoms. However, this was not the case (see Table 1). Additional control analysis, investigating whether changes in oscillatory activity are associated with MH and motor impairment (UPDRS-Part 3), did not show any significant frequency modulations in any brain region (for all interactions between MH and UPDRS-Part 3 the p-values > 0.05; FDR corrected; Supplementary Table S3) between groups as a function of patients’ motor impairment. Moreover, PD-MH and PD-nMH did not show any difference in the UPDRS motor scores (Table 1). These data show that the theta oscillatory changes associated with lower cognitive functions in PD-MH patients are not related to features of motor impairment.

### Lower center frequency (first assessment) is associated with lower cognitive frontal-subcortical cognitive functions in PD-MH patients, but not in PH-nMH patients

Previous research on cognitive impairment in PD has described a frequency “shift” or slowing down from predominant alpha oscillations to predominant theta oscillations, after the onset of dementia (for a review Aarsland et al., 2021). However, these interpretations were predominantly reached based on changes in power (intensity of the oscillation), rather than in the center frequency (the actual frequency - in Hz - at which the peak is observed). That is, the “shift” was inferred from heaving a reduced power in the alpha band and higher in the theta band, rather than a change in the frequency peak from 8-13Hz (alpha) to 4-8Hz (theta band). Therefore, we tested whether the association between theta oscillations, MH and lower frontal-cognitive functions is also associated with a change in the center frequency (shift of the center frequency in the 4-13Hz frequency band). We observed a significant (p-values <0.05; FDR corrected; Table S6) modulation of the center frequency in this theta-alpha range (Figure 3A) as a function of the interaction between the two terms (frontal-subcortical cognitive functions and MH (yes/no)). This interaction was localized over a central left electrode and a temporal electrode (Figure 3B and Supplementary Table S6). Post-hoc analyses show that for PD-MH the center frequency is significantly (p-value = 0.001, Figure 3C) associated with the frontal-subcortical cognitive functions, in which a lower center frequency (i.e. shifting from alpha to theta) is associated with lower cognitive functions. This association was not observed for PD-nMH (p-value = 0.49; Figure 3D). None of the other electrodes showed a significant interaction (all p-values > 0.05; FDR corrected; Supplementary Table S4). Additional analysis showed the absence of significant (all p-values > 0.05; FDR corrected; Supplementary Table S6) modulations of the center frequency for the posterior functions in the theta-alpha frequency range (interaction between the two terms (posterior functions and by MH (yes/no)). Collectively, these results show that the association between MH and frontal-subcortical cognitive function is due to both power and center frequency changes in the alpha-theta range (see supplementary results for center frequency changes associated with MH).

**Figure 3.**
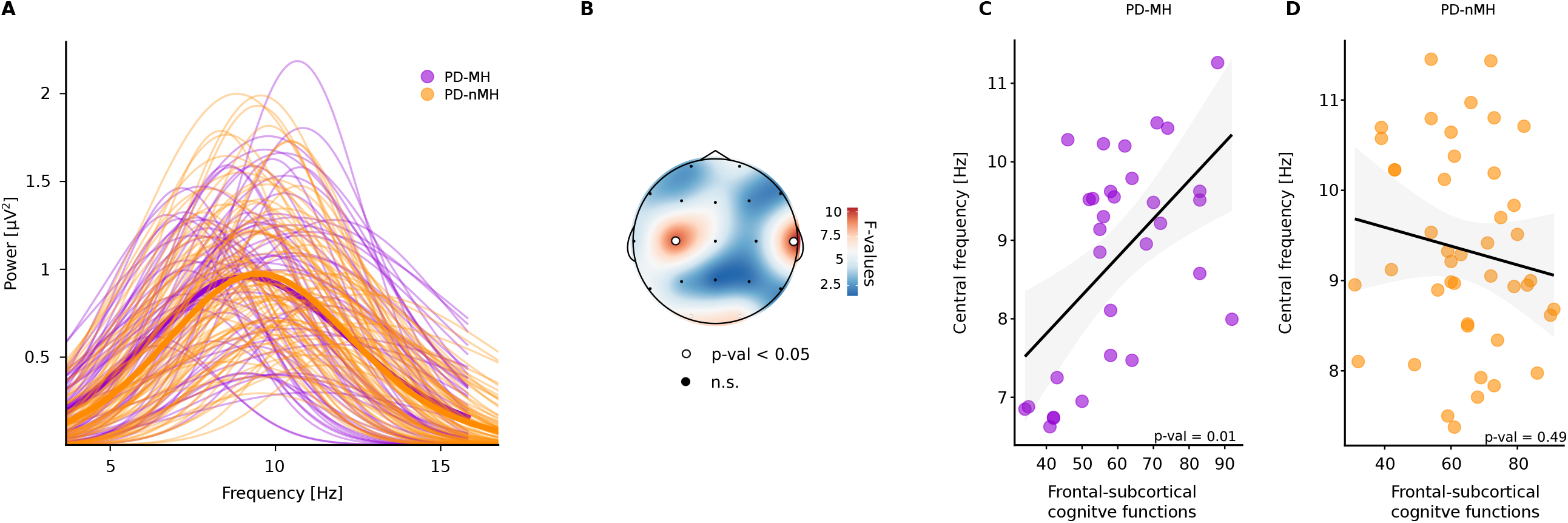
Association between frontal-subcortical cognitive functions and frontal central frequency in PD-MH. **A**. Reconstructed aperiodic-adjusted theta peaks, for PD-MH (violet) and PD-nMH (orange), thicker lines indicate the mean of each group. Thinner lines indicate the single patient data. **B**. Topography of F-values indicating the interaction between patients’ group and frontal-subcortical cognitive functions. White highlighted dots indicate electrodes showing a significant (p-values < 0.05; FDR-corrected) interaction for the center frequency (4-13Hz) and the frontal-subcortical cognitive functions. **C**. Center frequency is associated with frontal-subcortical cognitive functions in PD-MH. Lower center frequency is associated with lower cognitive functions. Single dots represent the value for each patient (average of the electrodes showing a significant interaction between patients’ group and frontal-subcortical cognitive functions). **D**. Center frequency is not associated with frontal-subcortical cognitive functions in PD-nMH. Single dots represent the value for each patient. Significance was obtained with permutation tests and multiple comparisons were corrected with FDR.

### Aperiodic signals (exponent and offset; first assessment) are not associated with cognitive functions

The precise neural mechanisms that alter the aperiodic exponent of intrinsic neural activity remain an active area of research. However, a flattened exponent of the aperiodic signal has been correlated with age (He et al., 2019) and with age-related cognitive decline (Tran et al., 2020; Voytek et al., 2015). Similarly, the offset of the aperiodic signal decreases with age (Cellier et al., 2021; Whitford et al., 2007). Therefore, we investigated whether modulations of exponent and/or offset might be related to changes as a function of MH (yes/no) and frontal-subcortical cognitive functions (interaction between the two terms). Our results show that neither exponent nor offset of the aperiodic signals are significantly modulated (all p-values > 0.05; Supplementary Table S7) as a function of cognitive function (frontal-subcortical nor posterior) and MH. These results demonstrate that the theta modulations we observed are due to a change in oscillatory power and not to aperiodic modulations.

### Second assessment (after 2 years) and third assessment (after 5 years) reveal that cognitive decline is more severe in PD-MH patients

Neuropsychological follow-up at the second assessment (2 years after the first assessment) was available in 67 patients and at the third assessment (5 years after the first assessment) in 53 patients of the total of 75 PD patients, included at the beginning of the study. Data show that the decline (over the 3 assessments) in frontal-subcortical cognitive functions was significantly different between the two groups of patients (p-value = 0.01; interaction between MH and follow-up; Figure 4; Supplementary Figure S6). Post-hoc analyses revealed that frontal-subcortical cognitive decline in PD-MH was significant when comparing the data from the first assessment with the third assessment (after five years) (p-value < 0.001; t(1,127) = 4.01), and when comparing the second with the third assessment (p-value < 0.01; t(1,126) = 3.3) (Figure 4A; Supplementary Table S8). This comparison was not significant when comparing data from the first assessment with the second assessment (after 2 years) (p-value = 0.41; t(1,125) = 0.83). For PD-nMH cognitive decline was not statistically different for either comparisons (i.e. neither when comparing the data from the first and second assessment (p-value = 0.16; t(1,126) = 1.42) nor from first and third assessment (p-value = 0.42; t(1,127) = 0.8)) (Figure 4B). Additional post-hoc analyses revealed that frontal-subcortical cognitive functions in PD-MH are lower than those in PD-nMH only at five year follow-up (p-value < 0.01; t(1,116) = -2.66) (Figure 4C). No statistical difference was observed at the second assessment (p-value = 0.45; t(1,102) = -0.76) or the first assessment (p-value = 0.33; t(1,96) = -0.97) (Figure 4A-B). Concerning posterior cognitive function, despite a reduction in these functions in both groups of patients (main effect of time: p-value < 0.01), no significant statistical difference was observed between the two sub-groups of patients (interaction: p-value = 0.39).

**Figure 4.**
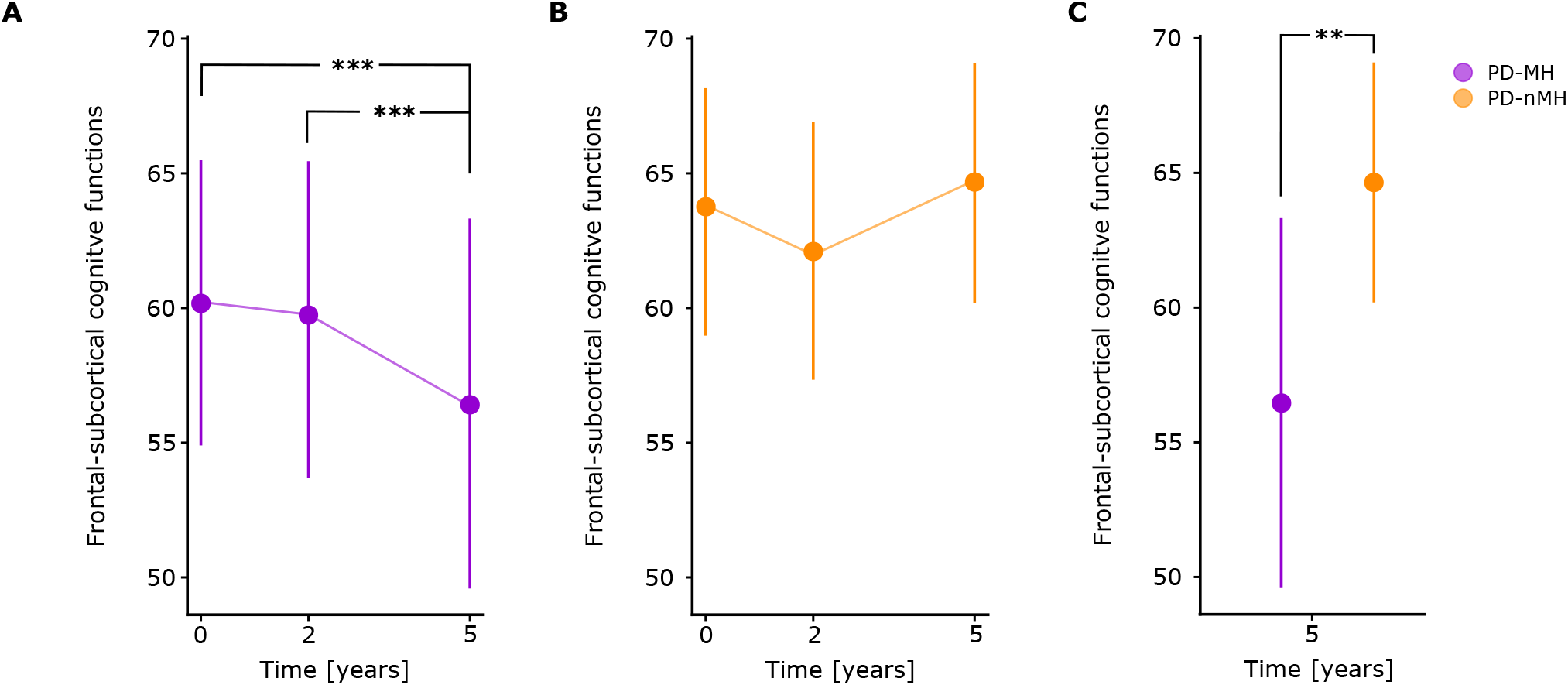
Longitudinal progression of the frontal-subcortical cognitive functions. **A**. In PD-MH patients frontal-subcortical cognitive functions decline significantly (p-value < 0.05) over the years. **B**. In PD-nMH patients frontal-subcortical cognitive functions do not decline significantly over 5 years. **C**. Difference in frontal-subcortical cognitive functions at the third assessment (year 5). PD-MH show a lower frontal-subcortical cognitive functions than PD-nMH. The bigger dots on the sides indicate the mean of the group. The error bars indicate 95% confidence interval. Asterisk indicates a statistical difference, it the p-value is less than 0.01, it is flagged with 2 asterisks (**), if the p-value is less than 0.001, it is flagged with three asterisks (***).

Collectively, these results show that the occurrence of MH in patients with PD is associated with a more severe form of the disease (without yet differing in the number of patients with MCI) characterized by a more important cognitive decline, especially for the frontal-subcortical cognitive functions.

### Cognitive decline in PD-MH patients (third assessment) is anticipated by the frontal-theta enhancement measured 5 years earlier (first assessment)

Informed by our results showing that both the frontal theta power and the theta-alpha center frequency (4-13Hz) are associated with frontal-subcortical cognitive score in patients with MH we investigated whether frontal theta power and/or the center frequency measured during the first assessment anticipated frontal-subcortical cognitive decline measured 5 years later. Concerning theta power, when assessing frontal-subcortical cognitive functions over 5 years, results show a significant interaction (p-value = 0.01) between the oscillatory power (measured during the first assessment) and the groups of patients. Post-doc analyses revealed that for PD-MH patients higher frontal theta power during the first assessment was associated with a stronger frontal-cognitive decline (p-value = 0.001; Figure 5A) 5 years later, which was not the case for patients without MH (p-value = 0.64; Figure 5B). None of the other frequencies showed this interaction (all p-value > 0.05; also see Supplementary results). Concerning the center frequency, we did not observe any significant association between this measure and the frontal-subcortical cognitive decline over 5 years (no interaction; p-value = 0.2). These results suggest that frontal theta power measured during the first assessment anticipates the cognitive decline occurring over 5 years, but only for PD-MH patients. In addition, these results show that by merging neuropsychological, neuropsychiatric, and EEG data, we are able to show that MH is associated with a higher risk of more rapid cognitive decline.

**Figure 5.**
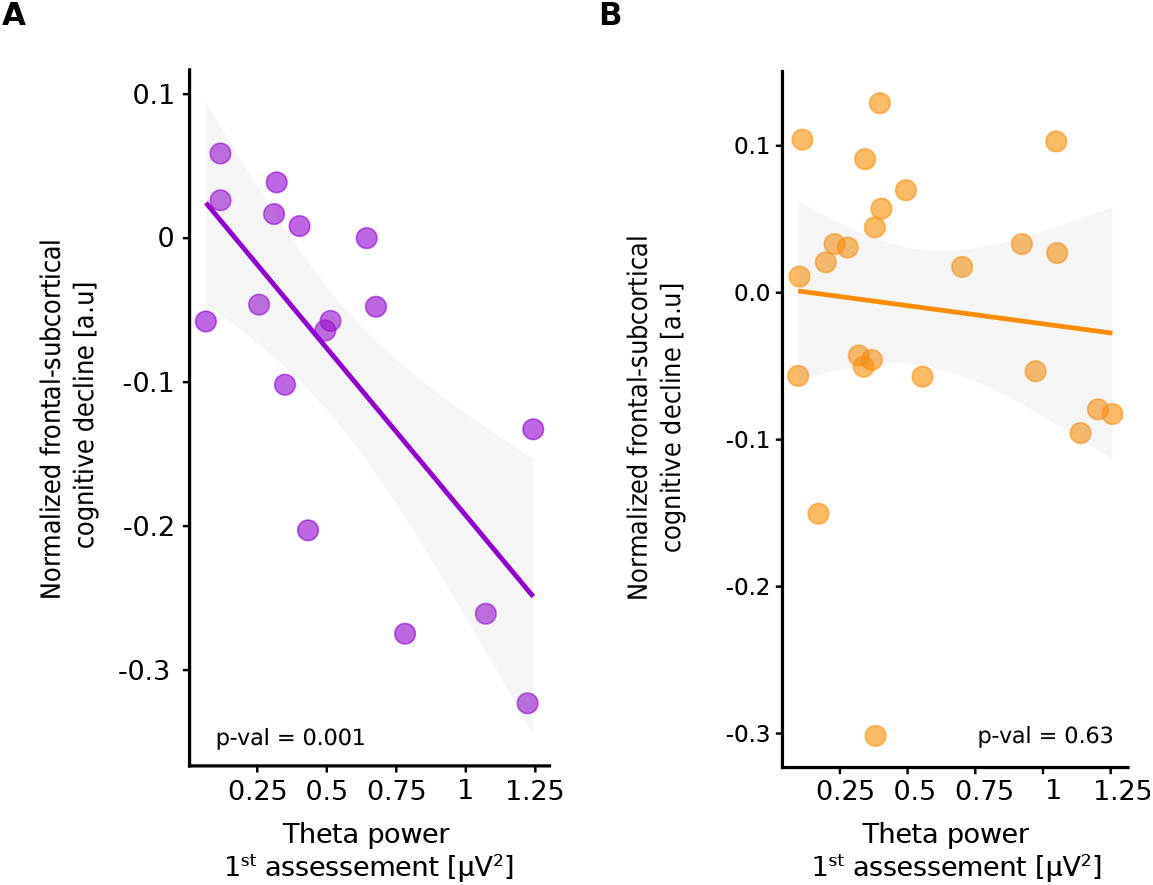
Frontal theta power during the first assessment anticipates cognitive decline occurring over 5 years. **A**. Results of the linear regression show that in PD-MH patients frontal theta power, as measured during the first assessment, is associated with the frontal-subcortical cognitive decline (normalized decline, see methods), as measured during the third assessment (5 years later). **B**. Results of the linear regression show that in PD-nMH patients frontal theta power, as measured during the first assessment, is not significantly associated with the frontal-subcortical cognitive decline (normalized decline, see methods), as measured during the third assessment (5 years later).

## Discussion

Conducting EEG, psychiatric and neuropsychological assessments in a group of 75 patients with PD, we report that specific alterations in frontal theta oscillations in PD patients with MH (Figure 2B, Figure 3B) were associated with a decrease in frontal-subcortical cognitive functions (Figure 2B-C, Figure 3B-C). Frontal theta oscillatory alterations were not associated with posterior cognitive functions and were absent in PD patients without MH (Figure 2D, Figure 3D). These data show that the combination of MH and enhanced theta oscillation allows to identify PD patients with a subclinical decrease in cognitive functions.

The detection of cognitive impairments in PD is of major clinical importance, as it is associated with earlier home placement, lower quality of life, and higher mortality (Aarsland et al., 2021; Chandler et al., 2021; Leroi et al., 2012). Cognitive impairment is a frequent non-motor symptom in PD, up to six times more common in PD than in the healthy age-matched population (Aarsland et al., 2001), and its prevalence increases with disease progression (Aarsland et al., 2021). Past neuropsychological and neuroimaging work in PD has identified deficits in posterior visual-perceptual, executive frontal-subcortical functions, and in related brain networks, thereby linking the neural mechanisms of hallucinations with cognitive impairments (Aarsland et al., 2003; Anang et al., 2014; Galvin et al., 2006; Uc et al., 2009). However, two main limitations have prevented the wider use of hallucinations as *early* predictors of cognitive decline in PD. First, previous work has focused on VH, which tend to occur in later stages of PD and hence in patients already showing advanced cognitive impairments, therefore, inevitably excluding them as an *early* marker of cognitive impairment. Second, although MH usually appear at earlier stages of the disease and before VH (Fénelon et al., 2000, 2011; Ffytche et al., 2017; Lenka et al., 2019), several clinical studies were not able to link them to cognitive dysfunction, failing to find significant cognitive impairments in PD patients with MH (Bejr-Kasem et al., 2021; Llebaria et al., 2010; Pagonabarraga et al., 2014). The present data provide evidence that MH allow to identify PD patients with cognitive impairment, however, only when MH are associated with specific EEG changes. Thus, when not taking EEG data into account, we confirm earlier data and show that PD patients with MH do not show stronger or distinct cognitive impairments, compared to PD patients without MH (Bejr-Kasem et al., 2021; Llebaria et al., 2010; Pagonabarraga et al., 2014). However, by combining psychiatric interviews and EEG data, we reveal that PD-MH patients with predominant theta (power and center frequency) oscillations have lower cognitive scores in frontal-subcortical functions (Figure 2C, Figure 3C). These findings are specific because frontal theta oscillatory alterations in PD-MH patients were not associated with changes in posterior cognitive functions and were absent in PD-nMH patients (Figure 2D, Figure 3D). Further control analyses revealed that MH-specific theta oscillations cannot be explained by differences in the occurrence of VH between groups, as only two of the investigated 75 patients reported structured VH, nor by any other of the clinical-demographic variables (e.g.age, disease duration, antiparkinsonian medication, or motor impairment), as the two groups of patients were similar in these variables.

Cognitive decline and dementia in PD, independently from hallucinations, have previously been associated with global (spatial and frequencies) EEG changes, characterized by enhanced theta oscillatory power as well as decreased power in higher frequency bands (i.e. alpha and beta). Comparable oscillatory abnormalities have also been reported in other neurodegenerative disorders characterized by cognitive decline, for instance, in dementia with Lewy bodies (Bonanni et al., 2008; Walker et al., 2015). However, previous results highlighting oscillatory changes were obtained by either comparing patients with different neurodegenerative diseases (e.g. Bonanni et al., 2008), by comparing PD patients with age-matched healthy controls, or by investigating PD patients at different stages of the disease and cognitive impairments (e.g. with vs. without dementia) (e.g. Hassan et al., 2017; Morita et al., 2011; Soikkeli et al., 1991; Stoffers et al., 2007; Wiesman et al., 2022). Accordingly, it is not known whether these previous oscillatory differences are only present in patients with cognitive deficits due to advanced PD, whether they depend on differences in disease duration or medication, or other clinical variables likely differing between previously tested groups. The present results extend and detail these findings. They are consistent with the importance of enhanced theta oscillations and cognitive decline in PD, but show changes that are more specific (as higher frequencies were not affected) and, critically, demonstrate them in a large group of patients with early PD and across two subgroups of patients that were clinically and demographically similar (differing only in MH), at a comparable stage of the disease, and were tested at a relatively early stage of the disease (mean of 5 years after PD diagnosis). The enhanced frontal theta activity in patients with cognitive decline that we described may reflect compensatory mechanisms caused by structural and functional changes (Buckner, 2004), reflecting disruption of thalamocortical circuits (Steriade et al., 1990) and/or differing involvement of prefrontal regions (i.e. Jensen et al., 2002; Raghavachari et al., 2001; Makeig et al., 2004; Bernasconi et al.,2021; Dhanis et al., 2021).

Moreover, we note that the present data on MH-specific theta alterations do not identify a general decrease in all tested cognitive functions and a global (whole-scalp) increase in theta oscillations. Instead, our analysis revealed a focal decrease in cognitive functions and a focal enhancement in theta oscillations. Compatible with proposals suggesting that reduced frontal dysfunction and frontal brain atrophy are associated with a higher risk for cognitive impairments in PD (Chung et al., 2019, 2020; Lee et al., 2014; Pagonabarraga et al., 2008), we report evidence that PD-MH patients show a decrease that is limited to frontal-subcortical functions and limited to an increase in frontal theta oscillations. This was corroborated by the absence of any differences in the tested posterior visual-perceptual functions (which are often observed in more advanced PD, associated with VH and dementia; Llebaria et al., 2010; Williams-Gray et al., 2007), by the absence of changes in posterior theta oscillations, and by the absence of changes in higher frequencies as well as aperiodic EEG signals. In conclusion, the present data suggest that a focal decrease in frontal-subcortical cognitive functions in early PD (i.e. 5 years after diagnosis and in stage 2 of the illness assessed by the Hoehn & Yahr scale) can de be identified by combining neuropsychological data with theta power over frontal-subcortical brain regions and the presence of MH.

The data from the 5-year longitudinal neuropsychological follow-up further corroborate and extend these findings, revealing that patients with MH have a stronger decline in frontal-executive functions (Figure 4) and that the enhanced frontal theta power, measured during the first assessment, is associated with a cognitive decline in frontal-subcortical functions occurring five years after the first assessment (Figure 5A). Frontal-subcortical deficits are believed to be an indicator of MCI (Pagonabarraga et al., 2008) and therefore predictor of later possible PDD. We note that the number of patients with MCI during the first assessment was not different between the two sub-groups, showing that MCI cannot explain the more prominent reduction in cognitive functions in the PD-MH group. Similarly, the frontal-subcortical cognitive functions during the first assessment were comparable between the two groups. This was different at the third assessment, carried out 5 years later, when PD-MH patients (versus PD-nMH patients) now showed a significant difference in in frontal-executive functions, with PD-MH heaving a significant more important cognitive impairment (the number of MCI patients was again not (yet) different between the two groups). Critically, our longitudinal data show that only frontal theta power anticipates the severity of the frontal-subcortical cognitive decline by 5 years and only in those patients reporting MH. Again, these EEG results were specific for frontal theta power as no other frequency anticipated the frontal cognitive decline. Compared to previous studies that suggested that enhanced theta oscillations (but also other variables) might indicate an increased risk of developing dementia (Cozac et al., 2016; Klassen et al., 2011), our data show that the combination of MH and selective frontal theta power allow a quantitative and early prediction of cognitive frontal-subcortical decline, an indicator for early MCI (Pagonabarraga et al., 2008). Finally, our results show that the theta power and not the center frequency in the theta-alpha band, anticipates cognitive decline in PD-MH patients and that the power changes are not due to changes in the background activity of the brain typically associated with aging (aperiodic signals; He et al., 2019; Voytek et al., 2015; Tran et al., 2020).

### Limitations of the study

Our findings should be considered in the context of the following limitations. First, although the use of an EEG system with a limited number of electrodes has several advantages for clinical use and patient comfort, a higher number of electrodes will allow for more extensive analyses (e.g. source localization). Second, we only measured EEG during the first assessment. Future studies should analyze frontal theta oscillations and other EEG signals also during follow-up examinations. Third, behavioral and imaging studies of cognitive tasks (e.g. Bejr-Kasem et al., 2022), in addition to resting-state EEG recordings, will be needed for a more in-depth characterization of the cognitive and neural impairments associated with MH.

Relatedly, we have recently developed a robotic procedure (Bernasconi et al., 2022) able to induce MH under robotically controlled conditions and further reported that PD-MH patients are highly sensitive to the procedure compared to healthy controls and PD-nMH patients (Bernasconi et al., 2021). Applying such methods will allow determining whether PD patients with heightened sensitivity to robot-induced MH can already be detected by enhanced frontal theta oscillations, enabling an even earlier identification of patients at risk to develop PD dementia, before MH and frontal-subcortical impairments become symptomatic.

## Methods

### Study design

Resting-state EEG data was collected using a 19-channel EEG system. Resting-state EEG data were collected with eyes open for a period lasting 5 minutes.

### Participants

Seventy-five individuals participated in the current study. All those fulfilling MDS new criteria for PD with minor hallucinations (PD-MH) — sense of presence, passage hallucinations, visual illusions and/or pareidolias (n = 31) — and without any hallucinations (PD-nMH; n = 44) were prospectively recruited from a sample of outpatients regularly attending to the Movement Disorders Clinic at Hospital de la Santa Creu i Sant Pau, Barcelona. Individuals were diagnosed with PD by a neurologist with expertise in movement disorders. Each individual was interviewed regarding disease onset, medication history, current medications, and dosage (levodopa daily dose and dopaminergic agonist-equivalent daily dose). Motor status and stage of illness were assessed by the MDS-UPDRS-III scale. The two subgroups of patients were comparable in gender, age, disease duration, dopaminergic doses, dopaminergic agonists, motor severity, sleep disturbances, and cognition.

Exclusion criteria were a history of major psychiatric disorders, cerebrovascular disease, conditions known to impair mental status other than PD, and the presence of factors that prevented MRI scanning (e.g. claustrophobia, MRI incompatible prosthesis). Patients with focal abnormalities in MRI or non-compensated systemic diseases (i.e. diabetes, hypertension) were also excluded. In patients with motor fluctuations, cognition was examined during the “on’’ state. All participants were on stable doses of dopaminergic drugs during the 4 weeks before inclusion. Patients were included if the hallucinations remained stable during the 3 months before inclusion in the study. No participant had used or was using antipsychotic medication. All subjects had a normal or corrected-to-normal vision. Informed consent to participate in the study was obtained from all participants according to the Declaration of Helsinki. The study was approved by the local ethics committee (#IIBSP-PAR-2019-17).

### Hallucinations and cognitive functions assessments

Minor hallucinations were assessed using the Hallucinations and Psychosis item of the MDS-UPDRS Part I and a semi-structured interview covering the types of minor hallucinations described in the literature. Patients were included in the PD-MH (i.e. with minor hallucinations) group if they had had a sense of presence (presence hallucinations), passage hallucinations, visual illusions and/or pareidolias at least monthly, and if these phenomena were present during the three months before inclusion in the study. Cognition was assessed by the Parkinson’s Disease-Cognitive Rating Scale (PD-CRS). To assess cognitive changes due to the disease and the presence of MH, cognitive functions were screened (PD-CRS) 24 and 60 months after the initial visit. During those follow-up assessments no EEG was recorded.

### EEG: acquisition preprocessing

Continuous EEG was acquired at 250 Hz from 19 standard scalp sites (Fp1-2, F3-4, C3-4, T3-4, T5-6, P3-4, O1-2, F7-8, Fz, Cz, Pz) using passive tin electrodes mounted in an elastic cap and referenced to the two mastoid leads. Vertical eye movements were monitored using a bipolar montage with two electrodes linked together and placed below each eye referenced to a third electrode placed centrally above the eyes. Horizontal eye movements were monitored using two electrodes placed on the external canthi of each eye. Electrode impedances were kept below 5 kOhm. The electrophysiological signals were filtered with a bandpass of 0.1–35 Hz and digitized at a rate of 250 Hz. EEG acquisition was done using the Brain Vision Recorder and BrainAMP system (Brain Products GmbH; Germany). Data were analyzed off-line with the EEGLAB toolbox for MATLAB (Delorme & Makeig, 2004; http://sccn.ucsd.edu/wiki/EEGLAB). After importing, data were low-pass-filtered at 45 Hz, and high-pass filtered at 1Hz with a conventional FIR filter. After removing the electrodes and epochs contaminated by artifacts (see below), data were re-referenced to the average reference.

The resting-state period was divided into 2-second epochs. This was done for each condition. Artifacts were removed in four steps: i) channels and then trials were rejected if their value exceeded 2.5 times the global variance (from all channels and trials, respectively). Frontal electrodes (Fp1 and Fp2) and EOG channels were excluded from this step to avoid biases (due to possible eye blinks) in the variance definition, and ii) independent component analysis (ICA) was applied to the remaining trials. ICA components reflecting eye blinks, saccades, or noise were identified and removed using SASICA (Chaumon et al., 2015). Noisy electrodes rejected at step i) were replaced by interpolating neighboring channels after the ICA procedure; iii) all epochs were inspected again for remaining ambient noise not removed by the ICA and rejected if artifacts had remained.

### Estimating oscillatory power and aperiodic signals

The power spectrum was estimated with the Fourier transform using Hanning as tapers, in the frequency 1Hz to 50Hz, with a frequency resolution of 0.5Hz. The power spectrum was calculated for each trial and then averaged. Power spectrum estimation was computed for each electrode and participant independently.

Electrophysiological data in humans are characterized by a prominent 1/f power distribution, often referred to as “background noise” (Bédard et al., 2006). This 1/f-like power distribution (also defined as aperiodic signal), captures the phenomenon whereby the power at low frequencies is greater, conversely power is progressively lower at higher frequencies. This pattern results in overall negatively sloped power spectrum across a wide range of frequencies. Most used methods and, therefore most of the research used to analyze oscillatory power neglect the role of this aperiodic signal. This introduces a possible confound, in which the identified neural oscillations contain a superposition of periodic and aperiodic signals. To avoid this confounding in our analyses we used a recent method that estimates both aperiodic and periodic signals (Donoghue et al., 2020).

We used the open-source, Python-based Fitting Oscillations and One-Over-F (FOOOF) (version 1.0) toolbox (Donoghue et al., 2020) to estimate both the periodic and aperiodic signals. We restricted the FOOOF algorithm to eight oscillatory peaks within the 2–45 Hertz range and constrained the peak width between 2-10Hz. We restricted the number of oscillatory peaks estimated by FOOOF to peaks, in order to reduce the risk of overfitting (see (Chiang et al., 2011; Dickson et al., 2010)). In addition, a threshold (a peak was greater than the noise floor of at least 1 standard deviation above the residuals) was used to label the peak as a genuine neural oscillation. All the other parameters were used as indicated by default (Donoghue et al., 2020). More details on the procedure can be found here (Donoghue et al., 2020). The Python code to fit the model was run in R Studio using the reticulate package (Ushey et al., 2020), and as implemented by (Ostlund et al., 2022). The model was applied to every electrode and patient.

## Statistical analysis

### Clinical-demographic variables

Statistical difference between PD-MH and PD-nMH on the measured clinical and demographic variables assessed using Welch test, Chi-squared, and U Mann Whitney test (see legend of Table 1 for details).

### Periodic and aperiodic EEG signals

To investigate modulations of the signals as a function of MH and cognitive functions models were performed with the periodic (i.e. power and center) or aperiodic signals (slope and offset) as the dependent variable, and with Group (PD-MH and PD-nMH) and cognitive functions (PDCRS; either for the frontal-subcortical sub-score or the posterior sub-score) as covariates (updrs-3 scores were used instead of the cognitive functions of the control analyses). An interaction term between the two covariates was used in the model. Models were independently applied to each electrode and frequency band of interest (theta, alpha, beta, gamma). The significance was estimated based on a permutation test for ANOVA (5000 iterations; permuco (Frossard & Renaud, 2021; R package). Correction for multiple comparisons for the number of electrodes was obtained with False Discovery Rate (FDR) (Benjamini & Yekutieli, 2001), applied to each frequency independently. Patients for which a peak in a determined frequency was not observed/measured were excluded from the statistical models for the given frequency band and/or electrode tested.

## Longitudinal data

### Progression of the cognitive functions

To assess changes in cognitive functions measured at baseline (Year 0), 24 months (Year 2) and 60 months (Year 5), frontal-subcortical and posterior cognitive functions were analyzed with linear mixed-effects models (Afex R package Singmann & Klauer, 2011)). Patients drop-out in the longitudinal data is due to either personal and/or clinical reasons. Two distinct models were performed for the two PD-CRS sub-scores. The models were performed with the cognitive scores and the measures time points as a fixed effect (interaction between the two terms) and with random intercepts for each participant. P-values were calculated using parametric bootstrap.

### EEG activity at baseline (Year 0) anticipates cognitive decline (at Year 5)

Based on the results of the cognitive decline over the year, frontal-subcortical cognitive decline was quantified by calculating the difference in score between the two points (Year 0 – Year 5) and normalized by dividing by the sum of the two scores (Year 0 + Year 5). To investigate whether the cognitive decline was associated with the theta power over the frontal electrodes (recorded at T0) and was differently modulated between subgroups of patients, we conducted permutation marginal tests for linear models (5000 iterations; *permuco* (Frossard & Renaud, 2021) R package), with theta (but also alpha, beta and gamma) power and subgroups as independent variables (and with an interaction term between the two), and cognitive decline as the dependent variable.

## Data availability

Please email all requests for academic use of clinical data to the corresponding authors. Requests will be evaluated based on institutional and departmental policies to determine whether the data requested is subject to intellectual property or patient privacy obligations. Data can only be shared for noncommercial academic purposes and will require a formal data use agreement.

## Code availability

Codes for the analyses are available here: https://gitlab.epfl.ch/fbernasc/pd_mh_eeg.git (will be made public after acceptance of the manuscript)

## Acknowledgements

We thank all patients for their participation to the study. We thank Prof. Andrea Serino and Prof. Gilles Allali for their comments on earlier version of the manuscript.

## Funding

This research was supported by two donors advised by CARIGEST SA (Fondazione Teofilo Rossi di Montelera e di Premuda and a second one wishing to remain anonymous) to O.B.; National Center of Competence in Research (NCCR) “Synapsy—The Synaptic Bases of Mental Diseases” grant number 51NF40-185897 to O.B.; Parkinson Suisse to O.B.; Bertarelli Foundation to O.B.; CIBERNED (Carlos III Institute) and FIS grant PI18/01717 to J.K.; Instituto de Salud Carlos III (ISCIII), Spain, to J.K.; PERIS, expedient number SLT008/18/00088 Generalitat de Catalunya to J. Pagonabarraga.

## Author contributions

F.B. analyzed the data and wrote paper, J.P., H.B-K, S.M-H collected data and wrote paper, J.K, O.B wrote paper. All authors approved the definitive version of the manuscript.

## Competing interests

All authors have no competing interests.

## Notes

### Competing Interest Statement

The authors have declared no competing interest.

