## Supplemental materials for "Theta oscillations and minor hallucinations in Parkinson’s disease reveal decrease in frontal lobe functions and later cognitive decline"

### *Supplementary methods*

*Periodic and Aperiodic EEG signals*

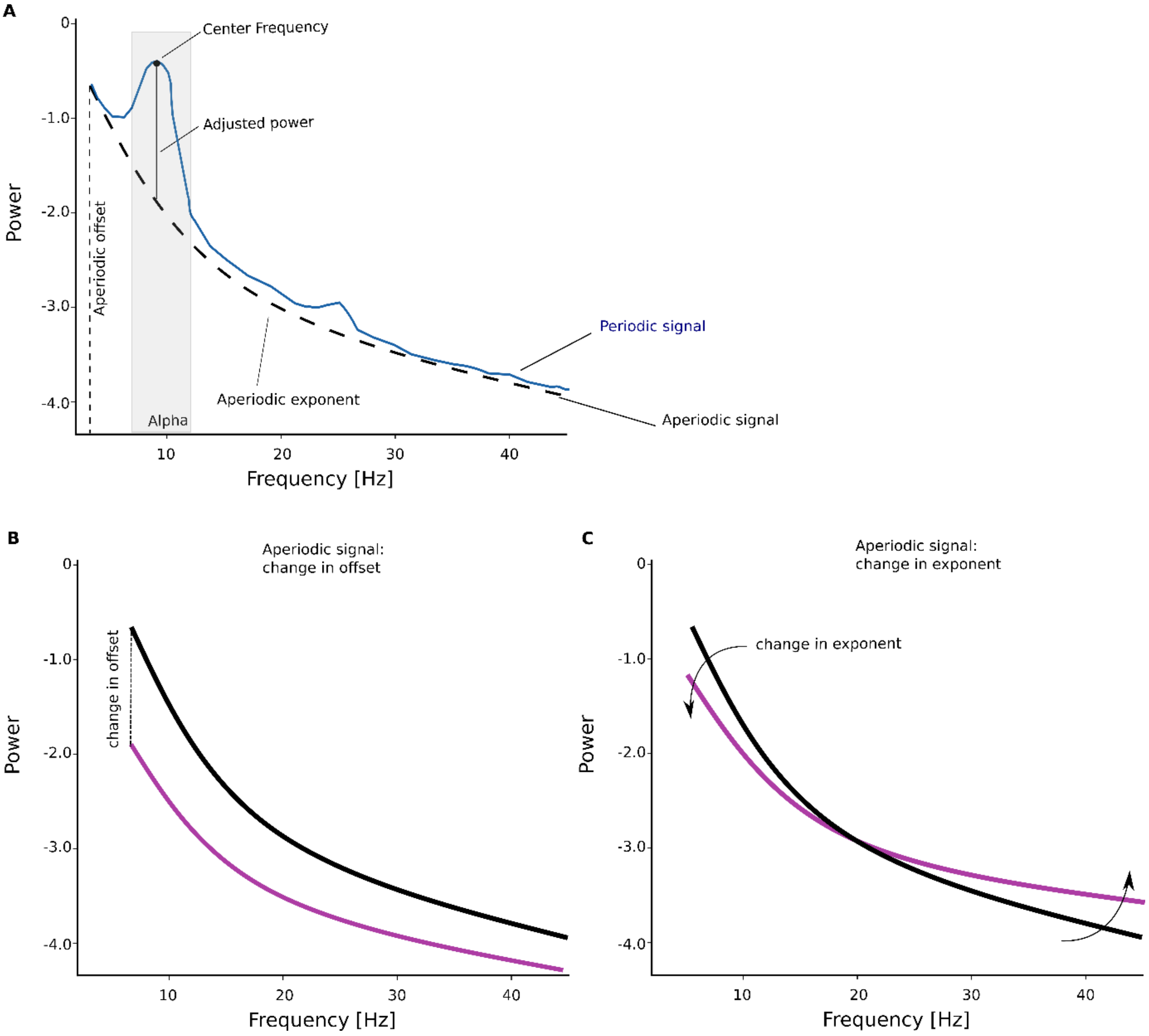


**Figure S1.** Illustration of a EEG signal. A. Illustration of a EEG power signal as two distinguishable signals: the aperiodic (1/f) background signal (dashed line); and the periodic components (dark blue) The periodic signal is composed of the: i) central frequency; ii) adjusted power; and iii) bandwidth (not shown). Those features of the periodic signal can change across groups and/or experimental conditions. The aperiodic signal can vary in Offset (or intercept) (B) or in exponent (or slope) (C).

***Supplementary results***

*Minor hallucinations (semi-structured interview)*

Presence hallucinations (PH) were the most predominant MH± present in 65% (20/31) of the patients± followed by passage hallucinations in 48% (15/31) and pareidolias in 35% (11/31) of patients (Figure S1). Our data further show that 26% (8/31) of the patients had exclusively PH; 23% (7/31) had concomitant PH and passage hallucinations; 16% (5/31) had only pareidolias or only passage hallucinations; 9.7% (3/31) concomitant PH and pareidolias; 6.5% (2/31) had concomitant PH; passage hallucinations and pareidolias; and 3.2% (1/31) had concomitant PSH and pareidolias.

*EEG model fitting: 'goodness of fit'*

The performance of the FOOOF algorithm was assessed via the 'goodness of fit' measures: $R^2$ and mean squared error (MAE), which represent the explained variance and total error of the model fit, respectively (Donoghue et al., 2020; Ostlund et al., 2022). These two parameters were estimated for each electrode and individual. Our results show that neither $R^2$ nor MAE are not significantly differing between patients PD-MH vs. PD-nMH in any electrodes (all p-values > 0.05; FDR corrected; Figure S2). There were 6 out of 1425 (0.42%) instances of underfitting (excluded from the analyses), defined as MAE > 0.100, when considering data from the full sample. There was no instance of potential overfitting (MAE = 0.199, R^2 = 0.996), defined here as MAE < 0.020. Collectively, these patterns indicate that the model fit parameters were appropriate across the examined frequency range. In addition, the absence of significant differences between the sub-groups of patients suggest that the results described are not due to a difference of the fitting procedure, but rather to a difference in the EEG features.

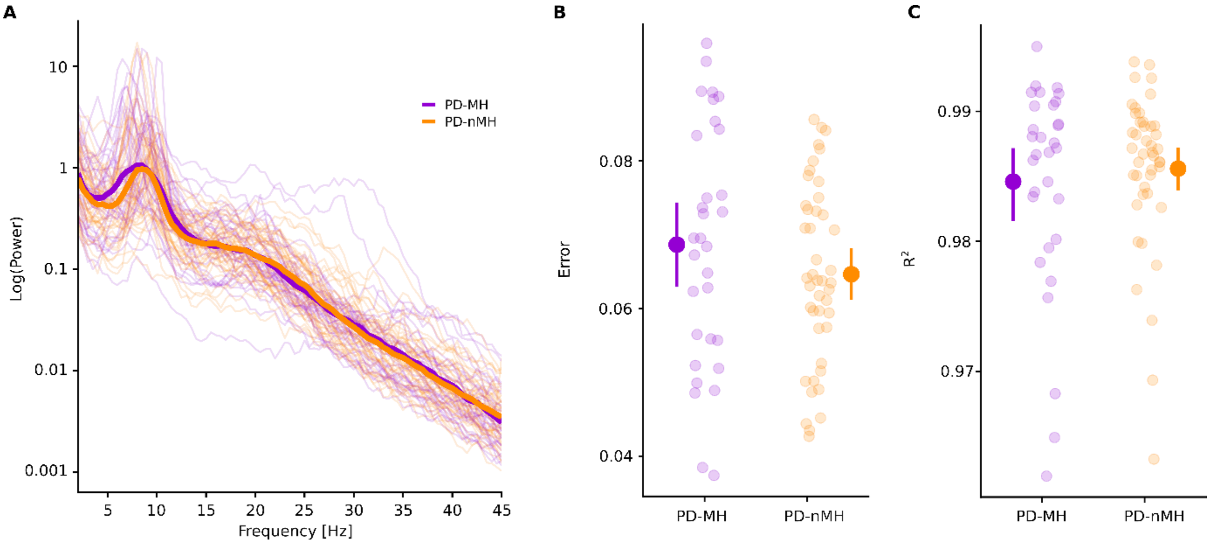


Figure S2. EEG raw power and fitting parameters. A. EEG power spectra for PD-MH (violet) and PD-nMH (orange), thicker lines indicate the mean of the groups, and the thinner lines indicate single patient data. B. R-squared values for the model fit for each patient for an exemplary electrode (F4). C. Error values for the model fit for each patient for an exemplary electrode (F4). The bigger dot on the side represents the mean and the error bars represent 95% confidence interval. No general differences in EEG frequency bands were observed between both patient groups.

***Frontal theta oscillatory power is enhanced in patients with MH and associated with frontal-subcortical cognitive functions (interaction between the two terms)***

| CHAN | EFFECT | F-VALUES | FREQ BAND | PERMUTATION P-VALUES (FDR-COR.) |
|---|---|---|---|---|
| FP1 | interaction | 0.05 | Theta | 0.87 |
| FP2 | interaction | 0.03 | Theta | 0.87 |
| F7 | interaction | 13.88 | Theta | 0.02* |
| F3 | interaction | 8.67 | Theta | 0.02* |
| FZ | interaction | 9.29 | Theta | 0.02* |
| F4 | interaction | 9.08 | Theta | 0.02 |
| F8 | interaction | 2.23 | Theta | 0.32 |
| T7 | interaction | 0.1 | Theta | 0.86 |
| C3 | interaction | 3.08 | Theta | 0.24 |
| CZ | interaction | 1.67 | Theta | 0.35 |
| C4 | interaction | 0.16 | Theta | 0.86 |
| T8 | interaction | 9.12 | Theta | 0.02* |
| P7 | interaction | 2.12 | Theta | 0.32 |
| P3 | interaction | 6.64 | Theta | 0.06 |
| PZ | interaction | 1.31 | Theta | 0.43 |
| P4 | interaction | 0.08 | Theta | 0.86 |
| P8 | interaction | 0.48 | Theta | 0.67 |
| O1 | interaction | 1.08 | Theta | 0.44 |
| O2 | interaction | 1.82 | Theta | 0.35 |
| FP1 | interaction | 0.3 | Alpha | 0.99 |
| FP2 | interaction | 0.45 | Alpha | 0.99 |
| F7 | interaction | 0 | Alpha | 0.99 |
| F3 | interaction | 0.12 | Alpha | 0.99 |
| FZ | interaction | 0.01 | Alpha | 0.99 |
| F4 | interaction | 0.02 | Alpha | 0.99 |
| F8 | interaction | 0.42 | Alpha | 0.99 |
| T7 | interaction | 0.5 | Alpha | 0.99 |
| C3 | interaction | 0.06 | Alpha | 0.99 |
| CZ | interaction | 0.33 | Alpha | 0.99 |
| C4 | interaction | 0.11 | Alpha | 0.99 |
| T8 | interaction | 0.28 | Alpha | 0.99 |
| P7 | interaction | 0 | Alpha | 0.99 |
| P3 | interaction | 0.17 | Alpha | 0.99 |
| PZ | interaction | 0.02 | Alpha | 0.99 |
| P4 | interaction | 0.23 | Alpha | 0.99 |
| P8 | interaction | 0.48 | Alpha | 0.99 |
| O1 | interaction | 0.09 | Alpha | 0.99 |
| O2 | interaction | 0.19 | Alpha | 0.99 |
| FP1 | interaction | 0.61 | Beta | 0.64 |
| FP2 | interaction | 0.92 | Beta | 0.64 |
| F7 | interaction | 0.65 | Beta | 0.64 |
| F3 | interaction | 1.29 | Beta | 0.64 |
| FZ | interaction | 0.02 | Beta | 0.9 |
| F4 | interaction | 0.05 | Beta | 0.86 |
| F8 | interaction | 0.09 | Beta | 0.86 |
| T7 | interaction | 0.63 | Beta | 0.64 |
| C3 | interaction | 0.28 | Beta | 0.75 |
| CZ | interaction | 2.76 | Beta | 0.64 |
| C4 | interaction | 2.25 | Beta | 0.64 |
| T8 | interaction | 0.14 | Beta | 0.85 |
| P7 | interaction | 0.37 | Beta | 0.75 |
| P3 | interaction | 1.6 | Beta | 0.64 |
| PZ | interaction | 1.82 | Beta | 0.64 |
| P4 | interaction | 0.88 | Beta | 0.64 |
| P8 | interaction | 0.63 | Beta | 0.64 |

| | | | | |
|---|---|---|---|---|
| O1 | interaction | 2.42 | Beta | 0.64 |
| O2 | interaction | 0.86 | Beta | 0.64 |
| FP1 | interaction | 0.61 | Gamma | 0.53 |
| FP2 | interaction | 0.81 | Gamma | 0.53 |
| F7 | interaction | 4.12 | Gamma | 0.18 |
| F3 | interaction | 2.94 | Gamma | 0.2 |
| FZ | interaction | 1.5 | Gamma | 0.35 |
| F4 | interaction | 0.07 | Gamma | 0.82 |
| F8 | interaction | 5.42 | Gamma | 0.18 |
| T7 | interaction | 4.14 | Gamma | 0.18 |
| C3 | interaction | 0.05 | Gamma | 0.82 |
| CZ | interaction | 3.78 | Gamma | 0.18 |
| C4 | interaction | 3.37 | Gamma | 0.19 |
| T8 | interaction | 2.44 | Gamma | 0.23 |
| P7 | interaction | 0.74 | Gamma | 0.53 |
| P3 | interaction | 2.33 | Gamma | 0.23 |
| PZ | interaction | 3.15 | Gamma | 0.19 |
| P4 | interaction | 0.62 | Gamma | 0.53 |
| P8 | interaction | 0.05 | Gamma | 0.82 |
| O1 | interaction | 3.69 | Gamma | 0.18 |
| O2 | interaction | 6.28 | Gamma | 0.18 |

**Table S1.** Statistical results of the models investigating modulations of the oscillatory power as a function of MH and frontal-subcortical cognitive functions (interaction between the two variables). Effect indicate which frequency is considered in the model. Permutation p-values are reported after FDR correction for multiple comparison.

***Oscillatory power is not modulated as a function of MH and posterior cognitive functions (interaction between the two terms)***

| CHAN | EFFECT | F-VALUES | FREQ BAND | PERMUTATION P-VALUES (FDR-COR.) |
|---|---|---|---|---|
| FP1 | interaction | 0.77 | Theta | 0.47 |
| FP2 | interaction | 0.12 | Theta | 0.77 |
| F7 | interaction | 6.44 | Theta | 0.1 |
| F3 | interaction | 4.78 | Theta | 0.13 |
| FZ | interaction | 6.26 | Theta | 0.1 |
| F4 | interaction | 6.47 | Theta | 0.1 |
| F8 | interaction | 3.38 | Theta | 0.17 |
| T7 | interaction | 0.35 | Theta | 0.63 |
| C3 | interaction | 0.06 | Theta | 0.82 |
| CZ | interaction | 2.95 | Theta | 0.19 |
| C4 | interaction | 1.83 | Theta | 0.31 |
| T8 | interaction | 4.23 | Theta | 0.15 |
| P7 | interaction | 2.81 | Theta | 0.19 |
| P3 | interaction | 0.84 | Theta | 0.47 |
| PZ | interaction | 1.21 | Theta | 0.39 |
| P4 | interaction | 0.45 | Theta | 0.6 |
| P8 | interaction | 1.76 | Theta | 0.31 |
| O1 | interaction | 3.39 | Theta | 0.17 |
| O2 | interaction | 5.94 | Theta | 0.1 |
| FP1 | interaction | 0 | Alpha | 0.99 |
| FP2 | interaction | 0.01 | Alpha | 0.99 |
| F7 | interaction | 0.17 | Alpha | 0.99 |
| F3 | interaction | 0.07 | Alpha | 0.99 |
| FZ | interaction | 0.6 | Alpha | 0.99 |
| F4 | interaction | 0 | Alpha | 0.99 |
| F8 | interaction | 1.66 | Alpha | 0.99 |
| T7 | interaction | 0.56 | Alpha | 0.99 |
| C3 | interaction | 1.13 | Alpha | 0.99 |
| CZ | interaction | 0.01 | Alpha | 0.99 |
| C4 | interaction | 0.71 | Alpha | 0.99 |
| T8 | interaction | 0.42 | Alpha | 0.99 |
| P7 | interaction | 0.9 | Alpha | 0.99 |
| P3 | interaction | 0 | Alpha | 0.99 |
| PZ | interaction | 0.08 | Alpha | 0.99 |
| P4 | interaction | 0.23 | Alpha | 0.99 |
| P8 | interaction | 0.69 | Alpha | 0.99 |
| O1 | interaction | 0.01 | Alpha | 0.99 |
| O2 | interaction | 0.11 | Alpha | 0.99 |
| FP1 | interaction | 0.09 | Beta | 0.94 |
| FP2 | interaction | 0.07 | Beta | 0.94 |
| F7 | interaction | 0.54 | Beta | 0.92 |
| F3 | interaction | 0.02 | Beta | 0.95 |
| FZ | interaction | 0.81 | Beta | 0.92 |
| F4 | interaction | 0.47 | Beta | 0.92 |
| F8 | interaction | 1.08 | Beta | 0.92 |
| T7 | interaction | 0.26 | Beta | 0.92 |
| C3 | interaction | 0.02 | Beta | 0.95 |
| CZ | interaction | 0.23 | Beta | 0.92 |
| C4 | interaction | 0.08 | Beta | 0.94 |
| T8 | interaction | 0.32 | Beta | 0.92 |
| P7 | interaction | 1.53 | Beta | 0.92 |
| P3 | interaction | 0.4 | Beta | 0.92 |
| PZ | interaction | 1.9 | Beta | 0.92 |
| P4 | interaction | 0.64 | Beta | 0.92 |

| | | | | |
|---|---|---|---|---|
| P8 | interaction | 1.01 | Beta | 0.92 |
| O1 | interaction | 0 | Beta | 0.97 |
| O2 | interaction | 0.52 | Beta | 0.92 |
| FP1 | interaction | 0.02 | Gamma | 0.89 |
| FP2 | interaction | 0.24 | Gamma | 0.75 |
| F7 | interaction | 1.05 | Gamma | 0.63 |
| F3 | interaction | 1.4 | Gamma | 0.63 |
| FZ | interaction | 1.4 | Gamma | 0.63 |
| F4 | interaction | 0.29 | Gamma | 0.75 |
| F8 | interaction | 0.28 | Gamma | 0.75 |
| T7 | interaction | 1.31 | Gamma | 0.63 |
| C3 | interaction | 0.47 | Gamma | 0.75 |
| CZ | interaction | 0.02 | Gamma | 0.89 |
| C4 | interaction | 1.29 | Gamma | 0.63 |
| T8 | interaction | 0.88 | Gamma | 0.63 |
| P7 | interaction | 0.16 | Gamma | 0.75 |
| P3 | interaction | 0.96 | Gamma | 0.63 |
| PZ | interaction | 1.44 | Gamma | 0.63 |
| P4 | interaction | 0.39 | Gamma | 0.75 |
| P8 | interaction | 1.41 | Gamma | 0.63 |
| O1 | interaction | 1.16 | Gamma | 0.63 |
| O2 | interaction | 0.3 | Gamma | 0.75 |

**Table S2.** Statistical results of the models investigating modulations of the oscillatory power as a function of MH and posterior cognitive functions (interaction between the two variables). Effect indicate which frequency is considered in the model. Permutation p-values are reported after FDR correction for multiple comparison.

***Oscillatory power is not modulated as a function of MH and motor impairment (UPDRS-part 3) (interaction between the two terms)***

| CHAN | EFFECT | F-VALUES | FREQ BAND | PERMUTATION P-VALUES (FDR-COR.) |
|---|---|---|---|---|
| FP1 | interaction | 0.08 | Theta | 0.95 |
| FP2 | interaction | 0.1 | Theta | 0.95 |
| F7 | interaction | 4.95 | Theta | 0.21 |
| F3 | interaction | 1.47 | Theta | 0.66 |
| FZ | interaction | 6.79 | Theta | 0.14 |
| F4 | interaction | 0.52 | Theta | 0.82 |
| F8 | interaction | 0 | Theta | 0.98 |
| T7 | interaction | 0 | Theta | 0.98 |
| C3 | interaction | 1.48 | Theta | 0.66 |
| CZ | interaction | 0.44 | Theta | 0.82 |
| C4 | interaction | 1.07 | Theta | 0.67 |
| T8 | interaction | 6.64 | Theta | 0.14 |
| P7 | interaction | 3.01 | Theta | 0.45 |
| P3 | interaction | 0.9 | Theta | 0.67 |
| PZ | interaction | 0.34 | Theta | 0.82 |
| P4 | interaction | 0.07 | Theta | 0.95 |
| P8 | interaction | 0.02 | Theta | 0.98 |
| O1 | interaction | 1.07 | Theta | 0.67 |
| O2 | interaction | 2.22 | Theta | 0.55 |
| FP1 | interaction | 1.06 | Alpha | 0.47 |
| FP2 | interaction | 1.55 | Alpha | 0.47 |
| F7 | interaction | 0.64 | Alpha | 0.51 |
| F3 | interaction | 1.47 | Alpha | 0.47 |
| FZ | interaction | 1.06 | Alpha | 0.47 |
| F4 | interaction | 0.62 | Alpha | 0.51 |
| F8 | interaction | 0.96 | Alpha | 0.47 |
| T7 | interaction | 1.6 | Alpha | 0.47 |
| C3 | interaction | 1.52 | Alpha | 0.47 |
| CZ | interaction | 3.63 | Alpha | 0.47 |
| C4 | interaction | 1.45 | Alpha | 0.47 |
| T8 | interaction | 2.35 | Alpha | 0.47 |
| P7 | interaction | 2.48 | Alpha | 0.47 |
| P3 | interaction | 0.23 | Alpha | 0.65 |
| PZ | interaction | 0.89 | Alpha | 0.47 |
| P4 | interaction | 1.07 | Alpha | 0.47 |
| P8 | interaction | 0.37 | Alpha | 0.6 |
| O1 | interaction | 1.07 | Alpha | 0.47 |
| O2 | interaction | 0.29 | Alpha | 0.63 |
| FP1 | interaction | 3.03 | Beta | 0.16 |
| FP2 | interaction | 1.21 | Beta | 0.29 |
| F7 | interaction | 2.75 | Beta | 0.16 |
| F3 | interaction | 3.73 | Beta | 0.16 |
| FZ | interaction | 2.91 | Beta | 0.16 |
| F4 | interaction | 1.43 | Beta | 0.28 |
| F8 | interaction | 4.23 | Beta | 0.16 |
| T7 | interaction | 1.24 | Beta | 0.29 |
| C3 | interaction | 2.44 | Beta | 0.17 |
| CZ | interaction | 5.06 | Beta | 0.16 |
| C4 | interaction | 3.61 | Beta | 0.16 |
| T8 | interaction | 2.75 | Beta | 0.16 |
| P7 | interaction | 3.67 | Beta | 0.16 |
| P3 | interaction | 3.36 | Beta | 0.16 |
| PZ | interaction | 3.64 | Beta | 0.16 |
| P4 | interaction | 0.5 | Beta | 0.47 |

| | | | | |
|---|---|---|---|---|
| P8 | interaction | 1.54 | Beta | 0.28 |
| O1 | interaction | 2.77 | Beta | 0.16 |
| O2 | interaction | 3.71 | Beta | 0.16 |
| FP1 | interaction | 0.52 | Gamma | 0.87 |
| FP2 | interaction | 0.08 | Gamma | 0.96 |
| F7 | interaction | 0.86 | Gamma | 0.71 |
| F3 | interaction | 1.18 | Gamma | 0.64 |
| FZ | interaction | 0.39 | Gamma | 0.87 |
| F4 | interaction | 0.03 | Gamma | 0.96 |
| F8 | interaction | 1.25 | Gamma | 0.64 |
| T7 | interaction | 3.71 | Gamma | 0.56 |
| C3 | interaction | 0.02 | Gamma | 0.96 |
| CZ | interaction | 3.64 | Gamma | 0.56 |
| C4 | interaction | 0.37 | Gamma | 0.87 |
| T8 | interaction | 1.85 | Gamma | 0.64 |
| P7 | interaction | 3.05 | Gamma | 0.56 |
| P3 | interaction | 0.01 | Gamma | 0.96 |
| PZ | interaction | 0 | Gamma | 0.96 |
| P4 | interaction | 0.28 | Gamma | 0.88 |
| P8 | interaction | 0.22 | Gamma | 0.88 |
| O1 | interaction | 1.73 | Gamma | 0.64 |
| O2 | interaction | 1.3 | Gamma | 0.64 |

**Table S3. S**tatistical results of the models investigating modulations of the oscillatory power as a function of MH and UPDRS-part 3 (interaction between the two variables). Effect indicate which frequency is considered in the model. Permutation p-values are reported after FDR correction for multiple comparison.

### *MH in PD are not associated with modulations of the oscillatory power*

Here, we investigated whether alterations in the oscillatory power was associated with MH. Our results show that, after correction for multiple comparisons, no electrodes showed significant (all p-values-values < 0.05; FDR corrected; Figure 3) modulation in oscillatory power as a function of MH (PD-MH vs. PD-nMH), in any of the tested frequency (see Table S5). It should be noted that when applying more complex models (i.e. to investigate power modulation as a function of MH and frontal-subcortical PD-CRS score), with an interaction between the two terms), frontal electrodes (F7-F3-Fz-F4-T8; same as those showing an interaction, Figure 2B) show a significant (p-values < 0.05; FDR corrected; main effect of MH) higher theta power for PD-MH (mean ± SD: 0.81 $\mu V^2$ ± 0.45 $\mu V^2$) than in PD-nMH (mean ± SD: 0.54 $\mu V^2$ ± 0.37 $\mu V^2$). However, because of the presence of an interaction (between MH and frontal-subcortical PD-CRS score) this main effect should be interpreted with caution.

Previous MEG research showed no difference in oscillatory activity between patients with complex hallucinations and patients without hallucinations. However, when comparing PD patients with only VH vs. patient without hallucinations data showed an enhanced theta oscillations (but also reduction in higher frequencies. This was observed for VH, over the whole scalp, and was further associated with a concomitant reduction of higher frequencies (Dauwan et al., 2019).

| CHAN | FREQUENCY | PERMUTATION P-VALUES (FDR-COR) | Power PD-MH MEAN ± SD | Power PD-nMH MEAN ± SD |
|---|---|---|---|---|
| C3 | Theta | 0.55 | 9.1 ± 1.43 | 9.56 ± 1.48 |
| C4 | Theta | 0.2 | 8.65 ± 1.31 | 9.51 ± 1.15 |
| CZ | Theta | 0.1 | 8.85 ± 1.53 | 9.41 ± 1.08 |
| F3 | Theta | 0.26 | 8.72 ± 1.52 | 9.1 ± 1.12 |
| F4 | Theta | 0.5 | 8.8 ± 1.62 | 9.07 ± 1.11 |
| F7 | Theta | 0.1 | 8.51 ± 1.34 | 9.16 ± 1 |
| F8 | Theta | 0.1 | 8.88 ± 1.26 | 9.46 ± 1.21 |
| FP1 | Theta | 0.1 | 8.53 ± 1.27 | 9.35 ± 1.09 |
| FP2 | Theta | 0.1 | 8.66 ± 1.39 | 9.39 ± 1.11 |
| FZ | Theta | 0.22 | 8.68 ± 1.52 | 9.2 ± 1.11 |
| O1 | Theta | 0.5 | 8.37 ± 1.37 | 8.99 ± 1.09 |
| O2 | Theta | 0.44 | 8.33 ± 1.32 | 8.87 ± 0.96 |
| P3 | Theta | 0.44 | 8.35 ± 1.12 | 8.91 ± 0.96 |
| P4 | Theta | 0.72 | 8.43 ± 1.18 | 9.09 ± 1.1 |
| P7 | Theta | 0.1 | 8.54 ± 1.5 | 9.31 ± 1.13 |
| P8 | Theta | 0.56 | 8.69 ± 1.52 | 9.01 ± 1.11 |
| PZ | Theta | 0.5 | 8.68 ± 1.43 | 9.14 ± 1.04 |
| T7 | Theta | 0.5 | 8.87 ± 1.61 | 9.49 ± 1.2 |
| T8 | Theta | 0.5 | 8.49 ± 1.4 | 9.12 ± 0.9 |
| C3 | Alpha | 0.99 | 0.96 ± 0.37 | 0.96 ± 0.26 |
| C4 | Alpha | 0.99 | 1.15 ± 0.49 | 1.2 ± 0.4 |
| CZ | Alpha | 0.99 | 1.15 ± 0.5 | 1.18 ± 0.41 |
| F3 | Alpha | 0.99 | 0.95 ± 0.44 | 0.94 ± 0.39 |
| F4 | Alpha | 0.99 | 0.93 ± 0.39 | 0.93 ± 0.31 |
| F7 | Alpha | 0.99 | 0.9 ± 0.47 | 0.92 ± 0.41 |
| F8 | Alpha | 0.99 | 1.14 ± 0.55 | 1.17 ± 0.4 |
| FP1 | Alpha | 0.99 | 0.86 ± 0.44 | 0.94 ± 0.41 |
| FP2 | Alpha | 0.99 | 0.87 ± 0.43 | 0.93 ± 0.44 |
| FZ | Alpha | 0.99 | 0.99 ± 0.43 | 0.99 ± 0.39 |
| O1 | Alpha | 0.99 | 0.99 ± 0.47 | 1.02 ± 0.3 |
| O2 | Alpha | 0.99 | 1.01 ± 0.49 | 1.11 ± 0.38 |
| P3 | Alpha | 0.99 | 0.76 ± 0.43 | 0.8 ± 0.38 |
| P4 | Alpha | 0.99 | 0.79 ± 0.34 | 0.8 ± 0.31 |

| | | | | |
|---|---|---|---|---|
| P7 | Alpha | 0.99 | 0.95 ± 0.59 | 1.08 ± 0.45 |
| P8 | Alpha | 0.99 | 0.76 ± 0.37 | 0.82 ± 0.26 |
| PZ | Alpha | 0.99 | 0.8 ± 0.44 | 0.76 ± 0.38 |
| T7 | Alpha | 0.99 | 0.93 ± 0.42 | 0.96 ± 0.3 |
| T8 | Alpha | 0.99 | 1 ± 0.57 | 1.12 ± 0.45 |
| C3 | Beta | 0.49 | 0.77 ± 0.29 | 0.83 ± 0.22 |
| C4 | Beta | 0.54 | 0.75 ± 0.27 | 0.81 ± 0.22 |
| CZ | Beta | 0.49 | 0.8 ± 0.33 | 0.88 ± 0.25 |
| F3 | Beta | 0.78 | 0.7 ± 0.31 | 0.73 ± 0.2 |
| F4 | Beta | 0.49 | 0.82 ± 0.25 | 0.91 ± 0.2 |
| F7 | Beta | 0.49 | 0.65 ± 0.27 | 0.72 ± 0.24 |
| F8 | Beta | 0.49 | 0.77 ± 0.3 | 0.84 ± 0.2 |
| FP1 | Beta | 0.49 | 0.56 ± 0.24 | 0.63 ± 0.2 |
| FP2 | Beta | 0.73 | 0.62 ± 0.25 | 0.65 ± 0.21 |
| FZ | Beta | 0.91 | 0.76 ± 0.3 | 0.77 ± 0.18 |
| O1 | Beta | 0.58 | 0.62 ± 0.3 | 0.67 ± 0.21 |
| O2 | Beta | 0.49 | 0.6 ± 0.26 | 0.67 ± 0.21 |
| P3 | Beta | 0.91 | 0.52 ± 0.22 | 0.52 ± 0.2 |
| P4 | Beta | 0.49 | 0.49 ± 0.2 | 0.55 ± 0.19 |
| P7 | Beta | 0.91 | 0.55 ± 0.26 | 0.56 ± 0.22 |
| P8 | Beta | 0.49 | 0.52 ± 0.17 | 0.58 ± 0.2 |
| PZ | Beta | 0.91 | 0.55 ± 0.24 | 0.55 ± 0.22 |
| T7 | Beta | 0.49 | 0.74 ± 0.28 | 0.83 ± 0.24 |
| T8 | Beta | 0.49 | 0.59 ± 0.28 | 0.65 ± 0.22 |
| C3 | Gamma | 0.93 | 0.2 ± 0.08 | 0.21 ± 0.09 |
| C4 | Gamma | 0.93 | 0.17 ± 0.08 | 0.21 ± 0.12 |
| CZ | Gamma | 0.93 | 0.21 ± 0.19 | 0.23 ± 0.12 |
| F3 | Gamma | 0.93 | 0.25 ±0.24 | 0.19 ± 0.14 |
| F4 | Gamma | 0.93 | 0.25 ± 0.18 | 0.25 ± 0.14 |
| F7 | Gamma | 0.93 | 0.2 ± 0.14 | 0.2 ± 0.16 |
| F8 | Gamma | 0.93 | 0.19 ± 0.11 | 0.22 ± 0.12 |
| FP1 | Gamma | 0.93 | 0.2 ± 0.13 | 0.19 ± 0.11 |
| FP2 | Gamma | 0.93 | 0.18 ± 0.09 | 0.2 ± 0.1 |
| FZ | Gamma | 0.93 | 0.28 ± 0.21 | 0.24 ± 0.21 |
| O1 | Gamma | 0.93 | 0.15 ± 0.11 | 0.17 ± 0.1 |
| O2 | Gamma | 0.93 | 0.17 ± 0.1 | 0.17 ± 0.08 |
| P3 | Gamma | 0.93 | 0.17 ± 0.09 | 0.16 ± 0.07 |
| P4 | Gamma | 0.93 | 0.14 ± 0.06 | 0.17 ± 0.07 |
| P7 | Gamma | 0.93 | 0.16 ± 0.09 | 0.15 ± 0.11 |
| P8 | Gamma | 0.93 | 0.18 ± 0.07 | 0.18 ± 0.09 |
| PZ | Gamma | 0.93 | 0.16 ± 0.1 | 0.16 ± 0.07 |
| T7 | Gamma | 0.93 | 0.2 ± 0.14 | 0.17 ± 0.09 |
| T8 | Gamma | 0.93 | 0.18 ± 0.12 | 0.16 ± 0.12 |

**Table S4.** Statistical results of the models investigating modulations of the oscillatory power for several frequency bands, as a function of MH. Statistical models were applied to each electrode independently. Permutation p-values are reported after FDR correction for multiple comparison. Results show that no statistical significant difference in oscillatory power is associated to MH.

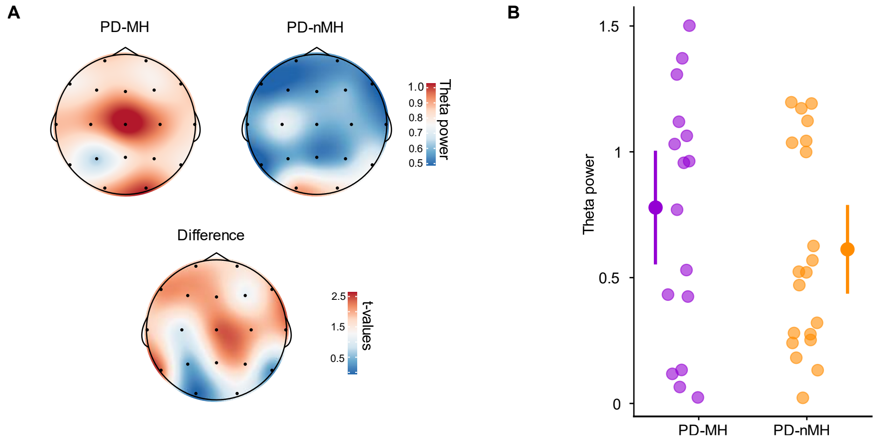


**Figure 3. Oscillatory theta power as a function of MH. A.** Topographies indicate the theta oscillatory power for PD-MH (left) and PD-nMH (right), and the topography on the bottom indicates the t-values for the statistical difference. No statistically significant difference was observed for oscillatory power between both patient groups. **B.** Oscillatory theta power for an exemplary electrode (F4). Single dots indicate the individual patient's theta power. The bigger dots on the sides indicate the mean of the group (PD-MH: violet; PD-nMH: orange). The error bars indicate 95% confidence interval.

### *MH in PD are associated with modulations of the center frequency in the theta-alpha frequency range*

Here, we investigated whether alterations in the theta-alpha center frequency were associated with MH. Our results show a significant change in center frequency in association with MH over fronto-central electrodes (p-values < 0.05; FDR corrected; Figure 4A). That is, PD-MH (mean ± SD: 8.59 ± 1.28 Hz) have a lower center frequency in the theta-alpha frequency band compared to PD-nMH (mean ± SD: 9.43 ± 1.12 Hz) (Figure 4B). The contribution of theta oscillations to hallucinations has been linked to several potential mechanisms such as enhanced theta-bursts in fronto-thalamic regions (Onofrj et al., 2019), and increased uncertainty in top-down activity resulting in incorrect sensory representations and hallucinations (i.e. Collerton et al., 2005). Further experimental-clinical work is needed to investigate whether the present findings on frontal theta oscillations link to experimentally-induced MH and related premotor and inferior frontal mechanisms (Bernasconi et al., 2021; Dhanis et al., 2021).

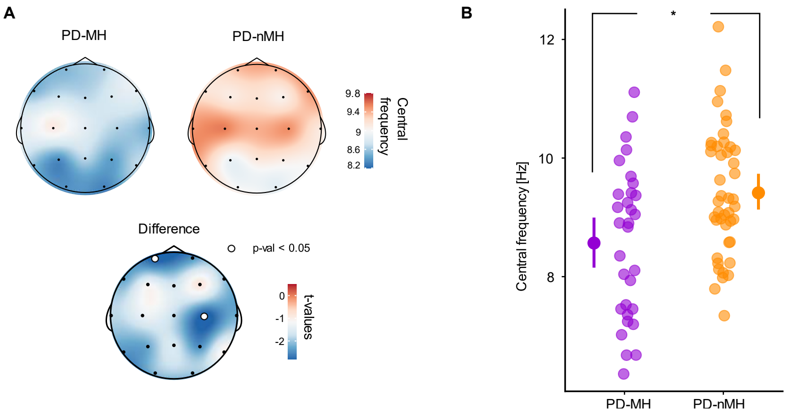


Figure 4. Center frequency as a function of MH . Topographies indicate the center frequency (in the 4-13Hz) for PD-MH (left) and PD-nMH (right), and the topography on the bottom indicates the t-values for the statistical difference. White highlighted dots indicated electrodes showing a significant interaction for the center frequency. D. PD-MH have a lower center frequency than PD-nMH. Single dots represent the value for each patient (average of the electrodes showing a significant interaction). The bigger dots on the sides indicate the mean of the group (PD-MH: violet; PD-nMH: orange). The error bars indicate 95% confidence interval. Asterisk indicates a significant difference.

| CHAN | FREQUENCY | PERMUTATION P-VALUES (FDR-COR) | Power PD-MH MEAN ± SD | Power PD-nMH MEAN ± SD |
|---|---|---|---|---|
| C3 | Theta-Alpha | 0.2 | 9.1 ± 1.43 | 9.56 ± 1.48 |
| C4 | Theta-Alpha | 0.049* | 8.65 ± 1.31 | 9.51 ± 1.15 |
| CZ | Theta-Alpha | 0.09 | 8.85 ± 1.53 | 9.41 ± 1.08 |
| F3 | Theta-Alpha | 0.24 | 8.72 ± 1.52 | 9.1 ± 1.12 |
| F4 | Theta-Alpha | 0.39 | 8.8 ± 1.62 | 9.07 ± 1.11 |
| F7 | Theta-Alpha | 0.05 | 8.51 ± 1.34 | 9.16 ± 1 |
| F8 | Theta-Alpha | 0.08 | 8.88 ± 1.26 | 9.46 ± 1.21 |
| FP1 | Theta-Alpha | 0.05 | 8.53 ± 1.27 | 9.35 ± 1.09 |
| FP2 | Theta-Alpha | 0.05 | 8.66 ± 1.39 | 9.39 ± 1.11 |
| FZ | Theta-Alpha | 0.12 | 8.68 ± 1.52 | 9.2 ± 1.11 |
| O1 | Theta-Alpha | 0.06 | 8.37 ± 1.37 | 8.99 ± 1.09 |
| O2 | Theta-Alpha | 0.08 | 8.33 ± 1.32 | 8.87 ± 0.96 |
| P3 | Theta-Alpha | 0.06 | 8.35 ± 1.12 | 8.91 ± 0.96 |
| P4 | Theta-Alpha | 0.05 | 8.43 ± 1.18 | 9.09 ± 1.1 |
| P7 | Theta-Alpha | 0.05 | 8.54 ± 1.5 | 9.31 ± 1.13 |
| P8 | Theta-Alpha | 0.3 | 8.69 ± 1.52 | 9.01 ± 1.11 |
| PZ | Theta-Alpha | 0.12 | 8.68 ± 1.43 | 9.14 ± 1.04 |
| T7 | Theta-Alpha | 0.09 | 8.87 ± 1.61 | 9.49 ± 1.2 |
| T8 | Theta-Alpha | 0.049* | 8.49 ± 1.4 | 9.12 ± 0.9 |

**Table S5.** Statistical results of the models investigating modulations of the center frequency in the theta-alpha (4-13Hz) frequency range, as a function of MH. Statistical models were applied to each electrode independently. Permutation p-values are reported after FDR correction for multiple comparison. Results show that PD-MH have a lower center frequency on central electrodes. Asterisks indicate significant effects.

***Center frequency is modulated as a function of MH and frontal-subcortical cognitive functions (but not with posterior cognitive functions)***

| CHAN | EFFECT | F-VALUES | COGNITIVE FUNCTIONS | PERMUTATION P-VALUES (FDR-COR.) |
|---|---|---|---|---|
| FP1 | interaction | 2.78 | Frontal-subcortical | 0.12 |
| FP2 | interaction | 5.28 | Frontal-subcortical | 0.06 |
| F7 | interaction | 3.52 | Frontal-subcortical | 0.11 |
| F3 | interaction | 4.13 | Frontal-subcortical | 0.09 |
| FZ | interaction | 5.22 | Frontal-subcortical | 0.06 |
| F4 | interaction | 2.42 | Frontal-subcortical | 0.14 |
| F8 | interaction | 2.25 | Frontal-subcortical | 0.15 |
| T7 | interaction | 4.44 | Frontal-subcortical | 0.09 |
| C3 | interaction | 9.54 | Frontal-subcortical | 0.03* |
| CZ | interaction | 5.33 | Frontal-subcortical | 0.06 |
| C4 | interaction | 3.42 | Frontal-subcortical | 0.11 |
| T8 | interaction | 10.34 | Frontal-subcortical | 0.02* |
| P7 | interaction | 5.65 | Frontal-subcortical | 0.06 |
| P3 | interaction | 3.21 | Frontal-subcortical | 0.11 |
| PZ | interaction | 1.53 | Frontal-subcortical | 0.21 |
| P4 | interaction | 1.88 | Frontal-subcortical | 0.18 |
| P8 | interaction | 2.52 | Frontal-subcortical | 0.14 |
| O1 | interaction | 6.54 | Frontal-subcortical | 0.06 |
| O2 | interaction | 6.98 | Frontal-subcortical | 0.06 |
| FP1 | interaction | 2.55 | Posterior | 0.24 |
| FP2 | interaction | 2.95 | Posterior | 0.24 |
| F7 | interaction | 1.82 | Posterior | 0.25 |
| F3 | interaction | 2.39 | Posterior | 0.24 |
| FZ | interaction | 3.32 | Posterior | 0.24 |
| F4 | interaction | 1.53 | Posterior | 0.25 |
| F8 | interaction | 1.05 | Posterior | 0.34 |
| T7 | interaction | 2.45 | Posterior | 0.24 |
| C3 | interaction | 3.86 | Posterior | 0.24 |
| CZ | interaction | 5.25 | Posterior | 0.24 |
| C4 | interaction | 2.53 | Posterior | 0.24 |
| T8 | interaction | 3.05 | Posterior | 0.24 |
| P7 | interaction | 3.13 | Posterior | 0.24 |
| P3 | interaction | 1.73 | Posterior | 0.25 |
| PZ | interaction | 0.11 | Posterior | 0.75 |
| P4 | interaction | 1.6 | Posterior | 0.25 |
| P8 | interaction | 1.5 | Posterior | 0.25 |
| O1 | interaction | 2.39 | Posterior | 0.24 |
| O2 | interaction | 1.65 | Posterior | 0.25 |

**Table S6. S**tatistical results of the models investigating modulations of the center frequency in the theta-alpha (4-13Hz) frequency band, as a function of MH and frontal-subcortical and posterior cognitive functions (interaction between the two variables; one model per cognitive functions). Effect indicate which cognitive function is considered in the model. Permutation p-values are reported after FDR correction for multiple comparison.

***Aperiodic signals are not significantly modulated by MH and frontal-subcortical cognitive functions (interaction between the two terms)***

Here, we investigated whether modulations of the aperiodic signal (Figure 5) might be related to changes as a function of MH (yes/no). Our results (Table S7) show that neither exponent (Figure 5A-B) nor the offset (Figure 5C-D) are significantly (all permutation p-values > 0.05; Supplementary Table S7) modulated by MH. These results demonstrate that the modulations we observed associated with MH (i.e. center frequency) are not due to aperiodic modulations.

| CHAN | APERIODIC SIGNAL | PERMUTATION P-VALUES (FDR-COR) | PD-MH MEAN ± SD | PD-nMH MEAN ± SD |
|---|---|---|---|---|
| C3 | Exponent | 0.89 | 1.55 ± 0.49 | 1.54 ± 0.38 |
| C4 | Exponent | 0.89 | 1.76 ± 0.29 | 1.69 ± 0.35 |
| CZ | Exponent | 0.89 | 1.75 ± 0.35 | 1.7 ± 0.38 |
| F3 | Exponent | 0.89 | 1.66 ± 0.4 | 1.62 ± 0.39 |
| F4 | Exponent | 0.89 | 1.76 ± 0.35 | 1.81 ± 0.27 |
| F7 | Exponent | 0.89 | 1.56 ± 0.42 | 1.6 ± 0.38 |
| F8 | Exponent | 0.89 | 1.88 ± 0.39 | 1.85 ± 0.33 |
| FP1 | Exponent | 0.89 | 1.56 ± 0.36 | 1.57 ± 0.36 |
| FP2 | Exponent | 0.89 | 1.58 ± 0.43 | 1.55 ± 0.4 |
| FZ | Exponent | 0.89 | 1.81 ± 0.36 | 1.77 ± 0.33 |
| O1 | Exponent | 0.89 | 1.57 ± 0.41 | 1.49 ± 0.4 |
| O2 | Exponent | 0.89 | 1.58 ± 0.35 | 1.5 ± 0.38 |
| P3 | Exponent | 0.89 | 1.52 ± 0.4 | 1.51 ± 0.37 |
| P4 | Exponent | 0.89 | 1.26 ± 0.55 | 1.23 ± 0.61 |
| P7 | Exponent | 0.89 | 1.51 ± 0.39 | 1.4 ± 0.41 |
| P8 | Exponent | 0.89 | 1.2 ± 0.63 | 1.36 ± 0.52 |
| PZ | Exponent | 0.89 | 1.56 ± 0.44 | 1.5 ± 0.42 |
| T7 | Exponent | 0.89 | 1.43 ± 0.55 | 1.5 ± 0.45 |
| T8 | Exponent | 0.89 | 1.59 ± 0.41 | 1.55 ± 0.47 |
| C3 | Offset | 0.83 | 0.51 ± 0.48 | 0.44 ± 0.44 |
| C4 | Offset | 0.83 | 0.75 ± 0.41 | 0.65 ± 0.47 |
| CZ | Offset | 0.83 | 0.73 ± 0.49 | 0.59 ± 0.47 |
| F3 | Offset | 0.83 | 0.67 ± 0.44 | 0.59 ± 0.44 |
| F4 | Offset | 0.86 | 0.71 ± 0.43 | 0.75 ± 0.4 |
| F7 | Offset | 0.97 | 0.59 ± 0.47 | 0.6 ± 0.41 |
| F8 | Offset | 0.83 | 0.8 ± 0.57 | 0.74 ± 0.43 |
| FP1 | Offset | 0.83 | 0.63 ± 0.44 | 0.54 ± 0.42 |
| FP2 | Offset | 0.83 | 0.65 ± 0.51 | 0.54 ± 0.44 |
| FZ | Offset | 0.83 | 0.77 ± 0.48 | 0.71 ± 0.44 |
| O1 | Offset | 0.83 | 0.77 ± 0.46 | 0.61 ± 0.46 |
| O2 | Offset | 0.83 | 0.75 ± 0.43 | 0.62 ± 0.47 |
| P3 | Offset | 0.83 | 0.71 ± 0.42 | 0.65 ± 0.5 |
| P4 | Offset | 0.83 | 0.51 ± 0.54 | 0.44 ± 0.6 |
| P7 | Offset | 0.83 | 0.76 ± 0.46 | 0.58 ± 0.46 |
| P8 | Offset | 0.97 | 0.44 ± 0.53 | 0.46 ± 0.48 |
| PZ | Offset | 0.83 | 0.73 ± 0.45 | 0.67 ± 0.41 |
| T7 | Offset | 0.97 | 0.37 ± 0.56 | 0.37 ± 0.46 |
| T8 | Offset | 0.83 | 0.82 ± 0.54 | 0.69 ± 0.58 |

**Table S7.** Statistical results of the models investigating modulations of the aperiodic signals (Offset and Exponent) as a function of MH and frontal-subcortical cognitive functions (interaction between the two variables). Effect indicate which between offset and slope are considered in the model. Permutation p-values (indicating the interaction term) are reported after FDR correction for multiple comparison.

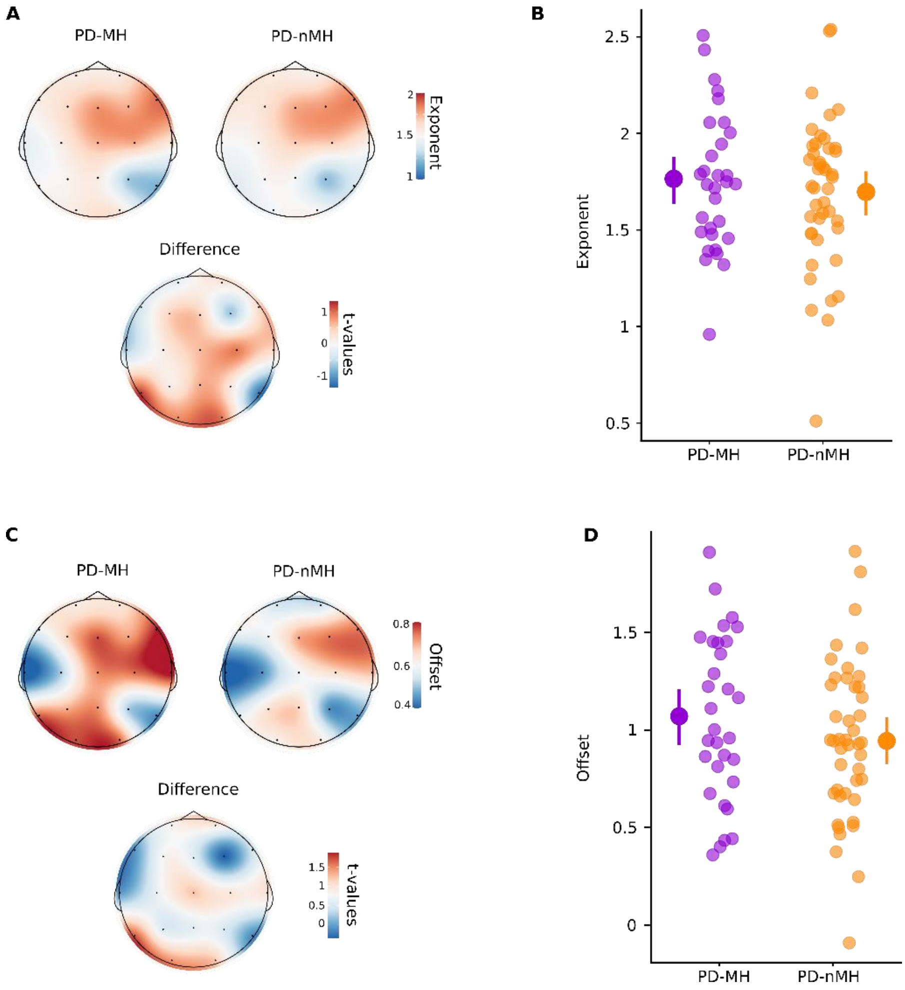


**Figure 5. Topographies showing aperiodic signals as a function of MH. A.** Topographies indicate the exponent (aperiodic signal) for PD-MH (left) and PD-nMH (right), and the topography on the bottom indicates the t-values for the statistical difference. No statistical difference was observed between patient groups. **B.** Offset for an exemplary electrode (Cz). Single dots indicate the individual participant's exponent. The bigger dots on the sides indicate the mean of the group (PD-MH: violet; PD-nMH: orange). The error bars indicate 95% confidence interval. **C.** Topographies indicate the exponent (aperiodic signal) for PD-MH (left) and PD-nMH (right), and the topography on the bottom indicates the t-values for the statistical difference. No statistical difference was observed between patient groups. **B.** Offset for an exemplary electrode (Cz). Single dots indicate the individual participant's offset. The bigger dots on the sides indicate the mean of the group (PD-MH: violet; PD-nMH: orange). The error bars indicate 95% confidence interval.

*Second assessment (after 2 years) and third assessment (after 5 years) reveal that cognitive decline is more severe in PD-MH patients)*

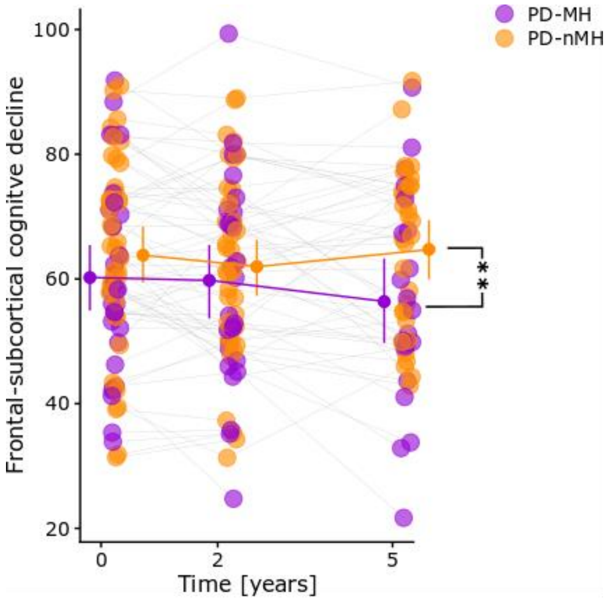


**Figure 6. Longitudinal follow-up of the frontal-subcortical cognitive functions.** Raw data for the five-year longitudinal PD-CRS follow-up data on the frontal-subcortical cognitive functions are shown. Single dots indicate an individual patient's cognitive score. The bigger dots on the sides indicate the mean of the group. The error bars indicate 95% confidence interval. Asterisk indicates a statistical difference.

| CONTRAST | ESTIMATE | SE | T-VALUES | P-VALUES |
|---|---|---|---|---|
| PD-MH TIME0 - PD-MH TIME2 | 1.51 | 1.82 | 0.83 | 0.41 |
| PD-MH TIME0 - PD-MH TIME5 | 8.17 | 2.02 | 4.05 | > 0.001* |
| PD-MH TIME2 - PD-MH TIME5 | 6.66 | 2.02 | 3.3 | > 0.01* |
| PD-NMH TIME0 - PD-NMH TIME2 | 2.23 | 1.58 | 1.41 | 0.16 |
| PD-NMH TIME0 - PD-NMH TIME5 | 1.37 | 1.7 | 0.8 | 0.42 |
| PD-NMH TIME2 - PD-NMH TIME5 | -0.87 | 1.73 | -0.5 | 0.62 |
| PD-MH TIME0 - PD-NMH TIME0 | -3.59 | 3.69 | -0.97 | 0.33 |
| PD-MH TIME2 - PD-NMH TIME2 | -2.87 | 3.76 | -0.76 | 0.45 |
| PD-MH TIME5 - PD-NMH TIME5 | -10.4 | 3.91 | -2.66 | 0.01* |

**Table S8.** Post-hoc comparisons of the frontal-subcortical cognitive functions, assessed longitudinally at two and five years, for both sub-groups of patients PD-MH and PD-nMH.

Additional analyses show that the two patient subgroups also not differ ($X^2$ (1,50) =0.16; p-value = 0.68) in the number of patients with mild cognitive impairment (defined as a PD-CRS total score ≤ 81 (Fernández de Bobadilla et al., 2013)), PD-MH (yes/no MCI): 7/15; PD-nMH (yes/no MCI): 8/22 neither after two years, nor after 5 years ($X^2$ (1,50) =1.46; p-value = 0.23), PD-MH (yes/no MCI): 7/15; PD-nMH (yes/no MCI): 8/22.

### *Cognitive decline in PD-MH patients (third assessment) is anticipated by the frontal-theta enhancement measured 5 years earlier (first assessment)*

To confirm the role of frontal theta oscillatory power during the first assessment for the anticipation of frontal-subcortical cognitive decline occurring over 5 years, we conducted an additional analyses, investigating whether the frontal theta would anticipate whether a patient has cognitive decline independently from MH. We separated the patients in two subgroup, depending on whether the normalized cognitive decline occurring over 5 years was lower than zero (i.e. with frontal-subcortical cognitive decline) and whether they had the cognitive decline equal to or bigger than zero. Our results show no significant (p-value = 0.29) interaction between the frontal theta and the groups (frontal-cognitive decline as dependent variable). Therefore, confirming the relevance of MH for identifying patients at higher risk of cognitive decline.

*List of references*


Bernasconi, F., Blondiaux, E., Potheegadoo, J., Stripeikyte, G., Pagonabarraga, J., Bejr-Kasem, H., Bassolino, M., Akselrod, M., Martinez-Horta, S., Sampedro, F., Hara, M., Horvath, J., Franza, M., Konik, S., Bereau, M., Ghika, J.-A., Burkhard, P. R., Ville, D. V. D., Faivre, N., … Blanke, O. (2021). Robot-induced hallucinations in Parkinson's disease depend on altered sensorimotor processing in fronto-temporal network. *Science Translational Medicine*, *13*(591). https://doi.org/10.1126/scitranslmed.abc8362

Collerton, D., Perry, E., & McKeith, I. (2005). Why people see things that are not there: A novel Perception and Attention Deficit model for recurrent complex visual hallucinations. *The Behavioral and Brain Sciences*, *28*(6), 737–757; discussion 757-794. https://doi.org/10.1017/S0140525X05000130

Dauwan, M., Hoff, J. I., Vriens, E. M., Hillebrand, A., Stam, C. J., & Sommer, I. E. (2019). Aberrant resting-state oscillatory brain activity in Parkinson's disease patients with visual hallucinations: An MEG source-space study. *NeuroImage: Clinical*, *22*, 101752. https://doi.org/10.1016/j.nicl.2019.101752

Dhanis, H., Blondiaux, E., Bolton, T., Faivre, N., Rognini, G., Van De Ville, D., & Blanke, O. (2021). Robotically-induced hallucination triggers subtle changes in brain network transitions. *NeuroImage*, *248*, 118862. https://doi.org/10.1016/j.neuroimage.2021.118862

Donoghue, T., Haller, M., Peterson, E. J., Varma, P., Sebastian, P., Gao, R., Noto, T., Lara, A. H., Wallis, J. D., Knight, R. T., Shestyuk, A., & Voytek, B. (2020). Parameterizing neural power spectra into periodic and aperiodic components. *Nature Neuroscience*, *23*(12), 1655–1665. https://doi.org/10.1038/s41593-020-00744-x

Fernández de Bobadilla, R., Pagonabarraga, J., Martínez-Horta, S., Pascual-Sedano, B., Campolongo, A., & Kulisevsky, J. (2013). Parkinson's disease-cognitive rating scale: Psychometrics for mild cognitive impairment. *Movement Disorders*, *28*(10), 1376–1383. https://doi.org/10.1002/mds.25568

Onofrj, M., Espay, A. J., Bonanni, L., Pizzi, S. D., & Sensi, S. L. (2019). Hallucinations, somatic-functional disorders of PD-DLB as expressions of thalamic dysfunction. *Movement Disorders*, *34*(8), 1100–1111. https://doi.org/10.1002/mds.27781

Ostlund, B., Donoghue, T., Anaya, B., Gunther, K. E., Karalunas, S. L., Voytek, B., & Pérez-Edgar, K. E. (2022). Spectral parameterization for studying neurodevelopment: How and why. *Developmental Cognitive Neuroscience*, 101073. https://doi.org/10.1016/j.dcn.2022.101073
